# Arabidopsis terminal nucleotidyl transferases govern secondary siRNA production at distinct steps

**DOI:** 10.1101/2024.05.27.596008

**Authors:** Maria Louisa Vigh, Axel Thieffry, Laura Arribas-Hernández, Peter Brodersen

## Abstract

In plants, RNA interference (RNAi) mediated by the endonucleolytic RNA-Induced Silencing Complex (RISC) defends against foreign RNA and regulates endogenous genes. Targeting of RISC to foreign RNA establishes amplification loops, wherein RNA-dependent RNA Polymerase 6 (RDR6) synthesizes double-stranded RNA (dsRNA) for secondary small interfering RNA (siRNA) biogenesis, using cleavage fragments of RNA targeted by RISC programmed with a primary siRNA as template. Secondary siRNA production from endogenous RISC targets requires a particular primary small RNA size or target site multiplicity. siRNA amplification in yeast and nematodes requires terminal nucleotidyl transferases (TNTases), but their roles in plants are unclear. Here, we demonstrate two functions of TNTases in siRNA amplification in *Arabidopsis thaliana*. URT1 prevents initiation of microRNA-induced secondary siRNA formation through uridylation of 5’-cleavage fragments, sometimes redundantly with the exosome and the TNTase HESO1. Once initiated via RDR6 recruitment, HESO1 and other TNTases stimulate secondary siRNA formation by producing 2-nt 3’overhangs on RDR6-synthesized dsRNA to yield substrates for processing into siRNAs by DICER-LIKE4. These results define molecular mechanisms by which TNTases control siRNA amplification in plants.

## INTRODUCTION

RNA interference (RNAi) phenomena rely on the RNA-Induced Silencing Complex (RISC) consisting of an ARGONAUTE (AGO) protein programmed with a small RNA of any sequence (Meister, 2013). RISC uses the small RNA as a guide to target RNA molecules sequence-specifically through base pairing. This process results in target RNA repression by a variety of mechanisms (Meister, 2013; Poulsen *et al*, 2013), sometimes via cleavage catalysed by the RNaseH-like activity of the AGO protein (Liu *et al*, 2004; Meister *et al*, 2004; Song *et al*, 2004). The likely ancestral role of RNAi is to eliminate parasitic genetic elements (Cerutti & Casas-Mollano, 2006), and many prokaryotes have minimal RNAi-like systems consisting only of AGO and a small RNA or DNA guide (Lisitskaya *et al*, 2018). Eukaryotic RNAi pathways involve dedicated small RNA processing systems, most of which rely on double-stranded RNA (dsRNA)-directed ribonucleases called Dicers for processing of dsRNA into small interfering RNA (siRNA) (Bernstein *et al*, 2001; Song & Rossi, 2017). Plants and animals have adopted RNAi independently for endogenous gene regulation through microRNAs (miRNAs) that are of fundamental importance for development and environmental adaptation (Axtell *et al*, 2011; Bartel, 2004).

Plants, fungi and many invertebrates encode a third core RNAi activity, RNA-dependent RNA Polymerase (RdRP) (Cerutti & Casas-Mollano, 2006), of key importance in the distinction between self and non-self RNA. In plants, siRNA populations targeting non-self RNA such as transgene- or virus-derived RNA are potently amplified by a system comprising the AGO protein AGO1, the RdRP RDR6, and its mandatory cofactors SUPPRESSOR-OF-GENE-SILENCING3 (SGS3) and SILENCING-DEFICIENT5 (SDE5), all central to antiviral RNAi (Dalmay *et al*, 2000; Fagard *et al*, 2000; Hernandez-Pinzon *et al*, 2007; Jauvion *et al*, 2010; Mourrain *et al*, 2000). siRNA amplification involves assembly of a complex of RISC, often containing AGO1, SGS3, SDE5 and RDR6 (Arribas-Hernandez *et al*, 2016; Sakurai *et al*, 2021; Yoshikawa *et al*, 2021; Yoshikawa *et al*, 2013) to direct dsRNA synthesis from a RISC target RNA. The first round of amplification relies on RISC loaded with what is sometimes referred to as a primary small RNA to emphasize the fact that its biogenesis does not depend on prior targeting by RISC; it may, for instance, be derived from direct processing of viral double-stranded RNA by the Dicer-like (DCL) enzyme DCL4 (Deleris *et al*, 2006). This first round of amplification gives rise to secondary siRNAs that may themselves engage in AGO1-RDR6-dependent reiterative amplification to establish a potent positive feedback system.

Several fundamental aspects of the plant siRNA amplification system remain incompletely understood. First, although many transgene- and virus-derived siRNAs and most endogenous miRNAs in the flowering plant *Arabidopsis thaliana* (Arabidopsis) associate with AGO1 and are 21 nucleotides (nt) in size, miRNAs generally do not trigger siRNA production by recruitment of the RDR6/SGS3/SDE5 amplifier module to their endogenous target RNAs (Howell *et al*, 2007). Second, plants have adopted the RDR6-dependent amplification module for endogenous gene regulation via phased or *trans*-acting siRNAs (Liu *et al*, 2020). The hallmark of this system is that the trigger miRNA never initiates unleashed positive feedbacks, but gives rise to a single round of secondary siRNA production, diagnosed by the 21-nt phase of siRNAs counting from the cleavage site guided by the trigger miRNA (Allen *et al*, 2005; Axtell *et al*, 2006; Liu *et al*., 2020). This property is of biological significance, because establishment of the positive feedback loop would irreversibly lock the target in the silent state, even in the absence of the original primary small RNA. The available evidence to explain these properties of the amplification system is largely descriptive but indicates clearly that one of two conditions must be met for miRNAs to trigger secondary siRNA production: Either the trigger miRNA is 22 nt in size (Chen *et al*, 2010; Cuperus *et al*, 2010), or the target RNA has more than one target site (“multi-hit target”), in which case the trigger miRNAs or siRNAs can be 21 nt in size (Axtell *et al*., 2006; Howell *et al*., 2007). In addition, a stalled ribosome juxtaposed to the miRNA target site strongly stimulates, and is sometimes required for, secondary siRNA formation (Iwakawa *et al*, 2021; Yoshikawa *et al*, 2016; Zhang *et al*, 2012). The facts that the secondary siRNAs made by DCL4 are 21 nt in size and that single-hit targets require 22-nt primary small RNAs to engage in secondary siRNA production conveniently explain why endogenous siRNA amplification does not lead to establishment of positive feedback loops. However, the more fundamental questions of how endogenous 21-nt miRNAs are prevented from triggering secondary siRNAs, which properties of 22-nt miRNAs overcome this barrier, and how establishment of reiterative AGO1-RDR6-dependent amplification through 21-nt siRNAs is confined to non-self RNA targets are not answered in a satisfactory way by these observations.

We previously showed that at least two endogenous activities are important for the limitation of miRNA-triggered secondary siRNA formation via RDR6 in Arabidopsis: the RNA exosome and the release factor-like mRNA quality control component PELOTA1 (PEL1) (Branscheid *et al*, 2015; Vigh *et al*, 2022). The exosome has a conserved role in degradation of 5’-cleavage fragments generated by RISC (Branscheid *et al*., 2015; Orban & Izaurralde, 2005) while the core biochemical function of PEL1 is to recognize stalled ribosomes and facilitate their resolution by catalysing subunit dissociation (Tsuboi *et al*, 2012). In Arabidopsis, these mutants have the important property that the strength of miRNA:target base pairing, not presence of stable RISC cleavage fragments, correlates with siRNA production (Branscheid *et al*., 2015; Vigh *et al*., 2022), arguing that RISC has a more direct involvement in RDR6 recruitment than mere production of cleavage fragments. In light of these observations, and the ability of catalytically inactive AGO1 programmed with a 22-nt miRNA to initiate secondary siRNA production (Arribas-Hernandez *et al*., 2016), we previously proposed a “RISC trigger model” in which long-lived RISC-target associations lead to recruitment of RDR6 (Fig 1A). The RISC trigger model predicts physical association between RDR6 and RISC specifically in situations that lead to amplified siRNA production (Arribas-Hernandez *et al*., 2016; Branscheid *et al*., 2015), as was later shown in elegant biochemical studies using a cell-free system that recapitulates miRNA-triggered secondary siRNA formation (Sakurai *et al*., 2021; Yoshikawa *et al*., 2021). The model also posits that acceleration of RISC-target RNA dissociation by exosome-mediated 5’-cleavage fragment degradation or by PEL1-dependent ribosome eviction underlies the requirement of these factors for limitation of secondary siRNA production. Nonetheless, only a subset of miRNAs become secondary siRNA triggers in exosome or *pel1* mutant backgrounds, suggesting that other systems may be used to counteract miRNA-triggered secondary siRNA production.

**Figure 1.**
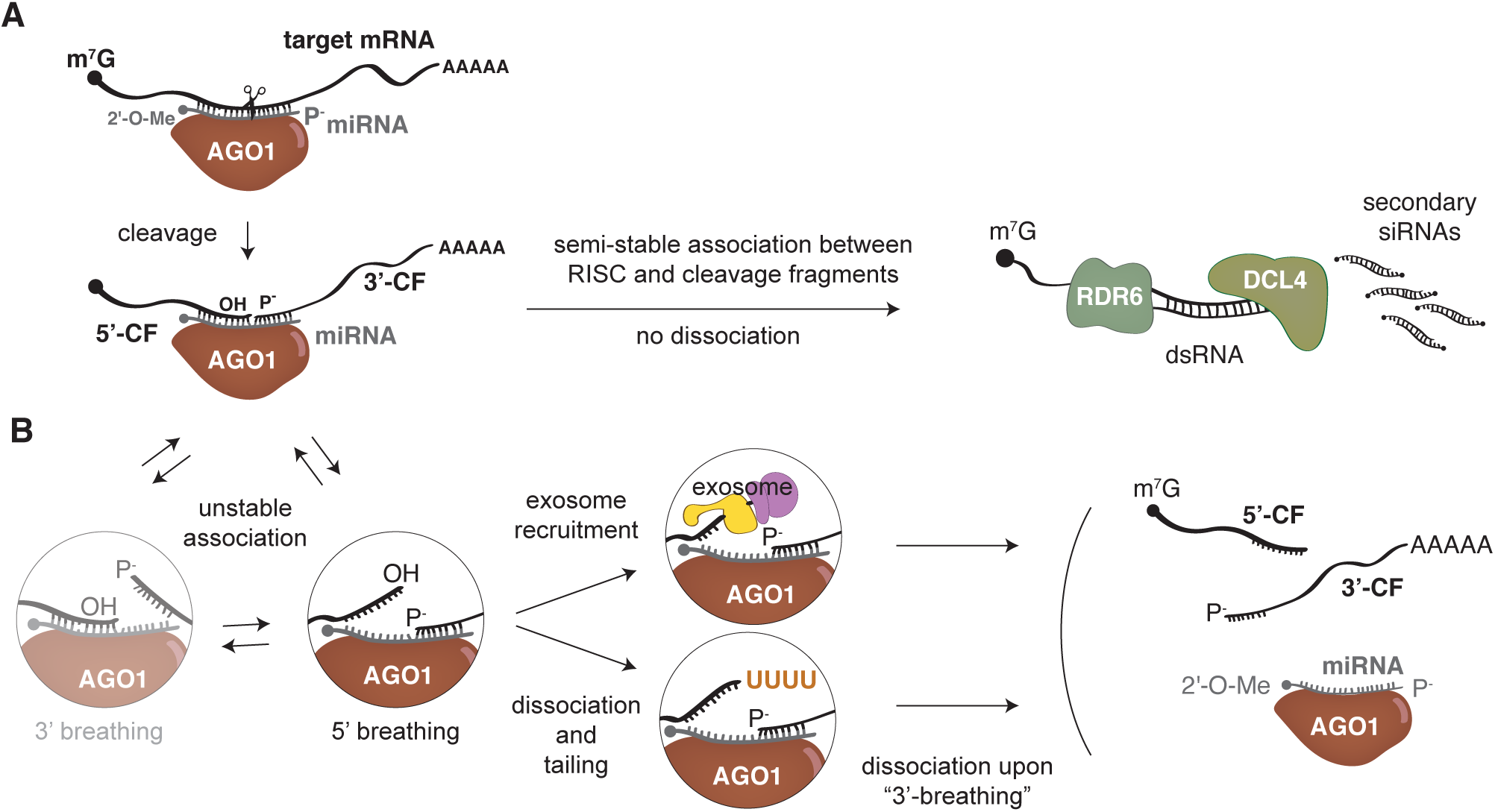
A hypothetical involvement of TUTases in limitation of secondary siRNA production. **A** In the RISC trigger model, a long-lived interaction between an AGO1-containing RISC and target RNA post cleavage leads to RDR6 recruitment, dsRNA synthesis and processing into secondary siRNAs by DCL4. **B** Two mechanisms may accelerate RISC dissociation post cleavage to limit secondary siRNA formation by targeting the 5’-cleavage fragment: Left, post cleavage, parts of the 5’- and 3’-cleavage fragments may dynamically breathe and reanneal with the AGO1-bound miRNA independently of one another. Right, upon breathing of the 5’-cleavage fragment, its 3’-end may be targeted for exosomal decay (top) or uridylation (bottom) preventing reannealing. In the absence of base pairing to the 5’-cleavage fragment, RISC dissociates upon breathing of the 3’-cleavage fragment before RDR6/SGS3/SDE5 recruitment.

Terminal nucleotidyl transferases (TNTases) add non-templated nucleotides to free 3’-ends of RNA. This versatile modification is linked to many aspects of small RNA biology. For example, uridylation of mammalian pre-miRNAs modulates Dicer activity, either optimizing it by extension to the preferred 2-nt 3’-overhang (Heo *et al*, 2012), or inhibiting it by synthesizing longer tails (Heo *et al*, 2009). Furthermore, in *Caenorhabditis elegans*, substrates for RdRPs may be marked by extending single-stranded RNAs with UG dinucleotides (Shukla *et al*, 2020), and in *Schizosacharomyces pombe*, the TNTase Cid12 is part of the RdRP complex and is necessary for siRNA amplification (Motamedi *et al*, 2004), albeit by an unknown mechanism. Finally, 5’-cleavage fragments produced by RISC-catalysed endonucleolysis are uridylated in many species, including plants (Shen & Goodman, 2004). Prominent enzymes responsible for such uridylation include the terminal uridyl transferases (TUTases) URT1 and HESO1 in Arabidopsis (Ren *et al*, 2014; Zuber *et al*, 2018), and current models propose that their uridylation directs cleavage fragment degradation (Ren *et al*., 2014). Nonetheless, the accumulation of 5’-cleavage fragments in *heso1* mutants is modest compared to the effect of mutation of the exosome or cytoplasmic exosome adaptors (Branscheid *et al*., 2015; Ren *et al*., 2014; Vigh *et al*., 2022), perhaps suggesting that uridylation of cleavage fragments may have other roles. For instance, one may envision that uridylation of 5’-cleavage fragments would impair reformation of base pairs upon “breathing” of duplexes between AGO-bound small RNA and the 5’-cleavage fragment in the RISC:target RNA complex post cleavage (Fig 1B). As such, uridylation may act to accelerate RISC dissociation from target mRNAs. In the RISC trigger model, such an effect is predicted to limit RDR6 recruitment and hence initiation of secondary siRNA formation by miRNA-programmed RISCs. Given this prediction we examined the roles of the two best characterized Arabidopsis TUTases, URT1 and HESO1 (De Almeida *et al*, 2018), in secondary siRNA formation. We show that URT1 reduces the initiation of secondary siRNA production from a small number of miRNA targets whose 5’-cleavage fragments undergo URT1-dependent uridylation. We also identify some cases in which URT1, HESO1 and the exosome collaborate to limit miRNA-induced secondary siRNA production. Interestingly, once secondary siRNA formation is initiated by RDR6 recruitment, HESO1 and additional unidentified TNTases promote the process by production of 2-nt 3’-overhangs optimal for processing into siRNAs by DCL4. Thus, TNTases are key players in secondary siRNA formation, and exert their roles at distinct steps in this pathway.

## RESULTS

### URT1 limits miRNA-triggered siRNA production from some target mRNAs

To test whether the TUTases URT1 and HESO1 have a role in limiting miRNA-triggered secondary siRNA production, we sequenced small RNA populations from mutants with T-DNA insertions in *URT1* and *HESO1*. We used the previously characterized *URT1* knockout allele *urt1-1* (Sement *et al*, 2013) and the *heso1-4* allele (GK-369H06) in which an exon encoding part of the nucleotidyl transferase domain is disrupted by the T-DNA insertion (Fig EV1A). We also included mutants in the cytoplasmic exosome cofactor *SKI2* (Branscheid *et al*., 2015) and in the core exosome component *RRP4* (Hematy *et al*, 2016) to obtain directly comparable results based on plant material, RNA and libraries produced and sequenced in parallel. Principal component analysis showed that the replicates clustered according to genotype, and that overall, the TUTase mutants had siRNA profiles distinct from exosome mutants and wild type (Fig 2A). We next conducted differential expression analysis of siRNAs specifically with an eye towards those produced from miRNA targets. The results showed that the *urt1-1* mutant had elevated levels of siRNAs from a small subset of miRNA targets distinct from those affected in exosome mutants (Figs 2B and EV1B). The clearest examples were found among targets that already give rise to secondary siRNAs in wild type plants, in particular targets of the 22-nt miRNAs miR393 (*AFB2*, *AFB3*, *TIR1*, *bHLH77*) (Howell *et al*., 2007) and miR472 (Boccara *et al*, 2014), and the multi-hit targets of the 21-nt *TAS1/2*-derived tasiRNAs and the 21-nt miRNAs miR161 and miR400 (Howell *et al*., 2007), exemplified by *RPF6* (Fig 2B,C and EV1C). siRNA production from the multi-hit targets was also increased in *ski2* and *rrp4* mutants, while miR393-triggered secondary siRNAs were only marginally affected by inactivation of *SKI2* or *RRP4* (Fig 2B,C and EV1C). Conversely, with the exception of miR161/miR400/tasiRNA-triggered siRNAs, the targets whose siRNA production was increased in exosome-related mutants were largely unaffected in the *urt1-1* mutant (Fig 2B,C and EV1C). The increased siRNA populations were localized adjacent to the cognate primary small RNA target sites (Fig 2C and EV1C), indicating that they were indeed triggered by RISC targeting. Importantly, plants carrying an independent knockout allele of *URT1* (*urt1-2* (Sement *et al*., 2013)) (Fig EV1A) showed an increase in *AFB2* siRNA production identical to that observed in *urt1-1* (Fig 2D), strongly supporting the conclusion that loss of *URT1* function causes increased *AFB2* siRNA production. This conclusion was further verified by restoration of low *AFB2* siRNA levels by transgenic expression of FLAG-tagged URT1 in the *urt1-1* knockout background (Fig 2E). We also verified that both *urt1-1* and *urt1-2* mutants expressed the miR393 trigger to levels similar to wild type (Fig 2D), excluding the trivial possibility that increased secondary siRNA levels might be a consequence of increased concentration of the miR393 trigger. Finally, we analysed whether the increased siRNA populations produced from miR393 targets resulted from illicit re-amplification triggered by 21-nt secondary siRNAs, or from more efficient 21-nt secondary siRNA production. These two scenarios can be distinguished by the phasing pattern of the *AFB2* siRNA population with respect to the miR393-guided cleavage site: illicit reamplification would erase, or greatly diminish, the phasing signature, while increased secondary siRNA production in the absence of reamplification would retain the phasing pattern seen in wild type. The results showed that the degree of phasing of the *AFB2* siRNA populations was similar in wild type and *urt1* mutants (Fig 2F). We conclude that *URT1* is required for limitation of secondary siRNA production from a subset of miRNA targets largely distinct from those affected by inactivation of the exosome. We note that the effect on secondary siRNA accumulation specifically is consistent with a role at the level of RDR6 recruitment as predicted by the RISC trigger model, but does not rule out effects on other steps, such as efficiency of dsRNA processing, or the rate of AGO loading or degradation of the resulting secondary siRNAs. We also note that in addition to showing defective limitation of secondary small RNA production from a subset of miRNA targets, our data confirm previous findings that inactivation of *URT1* leads to a broader production of ectopic siRNAs from non-miRNA targets transcriptome-wide (Fig EV1B) (Li *et al*, 2019; Scheer *et al*, 2021), probably linked to its other functions in mRNA uridylation, translational repression and decay (Scheer *et al*., 2021; Sement *et al*., 2013; Zuber *et al*, 2016).

**Figure 2.**
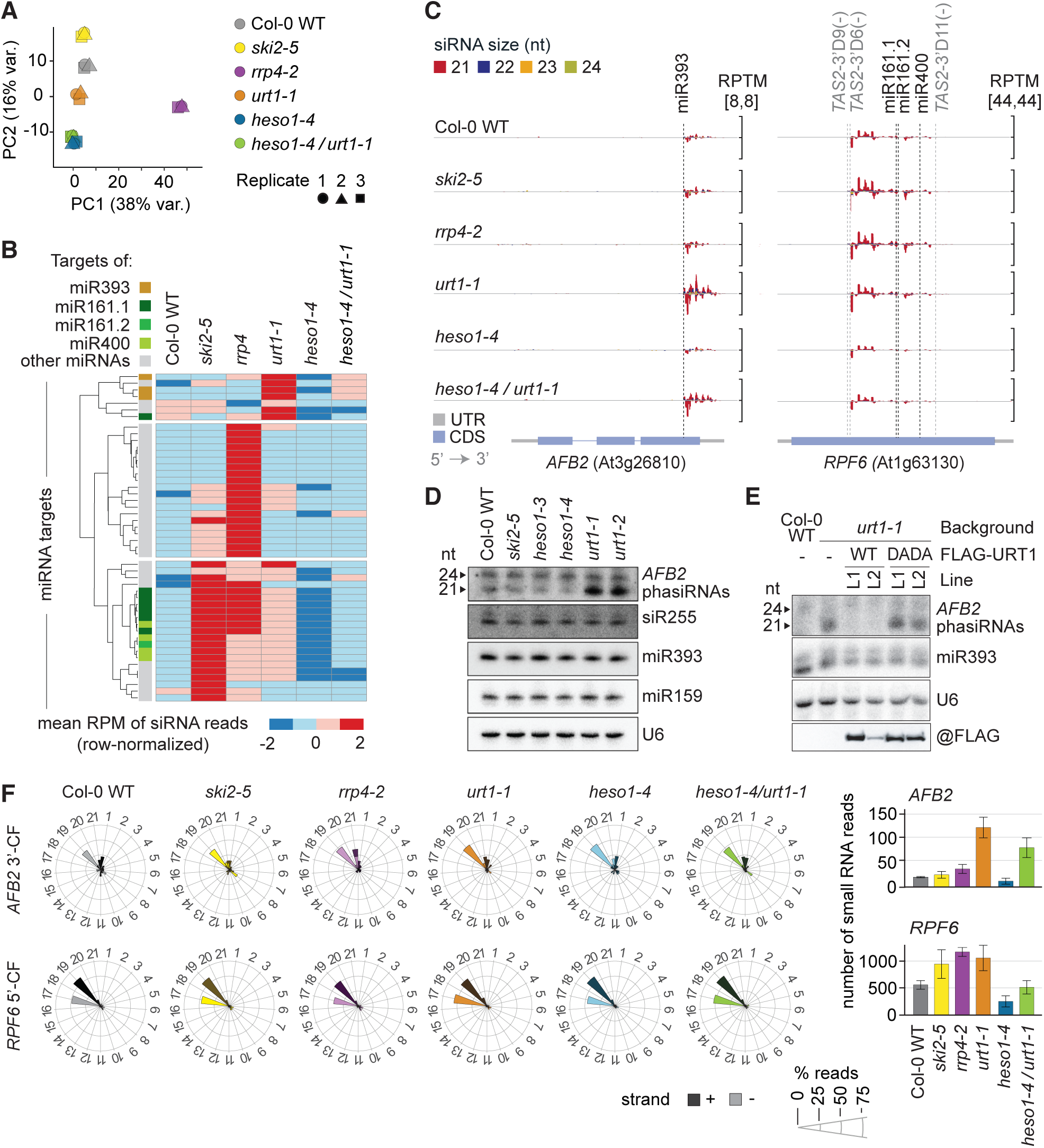
Effects of inactivation of the exosome, URT1 and HESO1 on miRNA-triggered secondary siRNA production. **A** Principal Component Analysis of the variance in the number of normalized mapped reads from small RNA libraries in three biological replicates of the indicated genotypes. **B** Heat map showing row-normalized z-scores of mean RPM (reads per million) of 21-nt siRNAs mapping to miRNA targets. Only targets with RPM>2 in at least one of the indicated genotypes, and with significantly different siRNA accumulation in at least one mutant compared to wild type (P_adj_ < 0.05) are shown. Information on differentially expressed siRNA populations is summarized in Dataset EV1. **C** 21-to 24-nt siRNAs mapping to *AFB2* and *RPF6* in the indicated genotypes. RPTM, reads per ten million. Reads above the X axes are in sense orientation, and below are antisense. Dashed lines indicate small RNA cleavage sites. Transcript architecture is represented below. **D** RNA blot analysis of total RNA from inflorescences of the indicated genotypes. Radiolabeled probes were hybridized consecutively to the same membrane. U6 is shown as loading control. **E** Top three panels, RNA blot analysis of total RNA extracted from seedlings of the indicated genotypes. Two independent transgenic lines of *urt1-1/FLAG-URT1* (WT) and *urt1-1/FLAG-URT1^DADA^* (DADA) were analyzed. Bottom panel, protein blot of FLAG-immunopurified fractions from 100 mg of seedling tissue. **F** Left, distribution of siRNAs mapping to *AFB2 and RPF6* among the 21 possible phases. Dark shades, reads mapping to the plus strand; light shades, reads mapping to the minus strand. Right, bar plots showing the total number of reads in each genotype. Error bars indicate standard error.

### Opposite effects of URT1 and HESO1 in miRNA-triggered secondary siRNA production

In contrast to *urt1-1*, the *heso1-4* knockout mutant did not show any instances of increased miRNA-triggered secondary siRNA production. Quite to the contrary, we observed several examples of miRNA targets with fewer siRNAs reads in *heso1-4* mutants than in wild type plants (Fig 2B). These examples included both the miR393-triggered *TIR1/AFB2/AFB3* siRNA populations analysed in the context of *URT1* mutations and several *TAS1/2*-miR161-miR400 multiple-hit triggered siRNAs mapping to pentatricopeptide-repeat protein (PPR)-encoding mRNAs (Figs 2B,C and EV1C). The effect of inactivating *HESO1* was the same in wild type and *urt1-1* genetic backgrounds, seen most clearly for miR393 targets (Figs 2C and EV1C). Thus, the double mutants did not reveal obvious epistatic relationships between the *urt1-1* and *heso1-4* mutations, precluding clear interpretations on relative functions of the two proteins in secondary siRNA production in a wild type background. The remainder of this report focuses on understanding the effects of the two TUTases on miRNA-triggered secondary siRNA production observed in this initial sequencing experiment. We treat the URT1-mediated limitation of miRNA-triggered secondary siRNA production first and return to the requirement for HESO1 for fully efficient secondary siRNA production subsequently.

### URT1-mediated limitation of AFB2-derived siRNAs requires its catalytic activity

The hypothesis that the uridylation of 5’-cleavage fragments is responsible for limitation of secondary siRNA production makes at least two additional testable predictions. (i) Point mutants specifically defective in catalytic activity of URT1 should show the same effect on miRNA-induced siRNA production as knockout mutants, and (ii) 5’-cleavage fragments of miRNA targets with increased siRNA production in *urt1* mutants should exhibit URT1-dependent uridylation. To test the first prediction, we used the *urt1-1* knockout background to express FLAG-tagged wild type URT1, and URT1 containing two point mutations (D491A/D493A, “DADA”) in aspartate residues required for Mg^II^ coordination in the catalytic center. The URT1^DADA^ mutant has no nucleotidyl transferase activity *in vitro* (Tu *et al*, 2015b). In contrast to URT1^WT^, URT1^DADA^ failed to complement the defect in limiting *AFB2* siRNA production of *urt1-1* (Fig 2E). We verified that the expression level of the miR393 trigger was unaffected, and that the protein levels of URT1^WT^ and URT1^DADA^ were similar in independent, stable transgenic lines (Fig 2E). Thus, the catalytic activity of URT1 is required for limitation of *AFB2* siRNA production.

### URT1 mediates uridylation of AFB2 5’-cleavage fragments

We next used 3’-RACE-seq to characterize the 3’-end of *AFB2* 5’-cleavage fragments. Because uridylation might involve multiple TUTases, and because 3’-tailed RNAs may display accelerated degradation via the exosome (Ibrahim *et al*, 2006), we completed this analysis in several mutant backgrounds to increase the likelihood of detecting U-tailed species. Specifically, we analysed wild type vs. *urt1-1*, and *heso1-4* vs. *heso1-4/urt1-1* to detect uridylation and its potential URT1 dependence. We also included the *rrp4-2* mutant with a partial loss-of-function mutation in a core exosome subunit (Hematy *et al*., 2016) to test whether uridylated cleavage fragments may be more abundant in this background, as exosomal degradation of nucleotidylated RISC 5’-cleavage fragments is suggested by previous analyses in both Arabidopsis and *Chlamydomonas reinhardtii* (Branscheid *et al*., 2015; Ibrahim *et al*., 2006). The 5’-cleavage fragment of the miR159 target *MYB33* was included as a positive control, because previous 3’-RACE-seq analyses have shown hierarchical uridylation of this fragment by HESO1 and URT1 (Zuber *et al*., 2018). Indeed, our analyses clearly detected HESO1-dependent U-tails and a minor, additive effect of *URT1* and *HESO1* mutations on *MYB33* 5’-cleavage fragments (Fig 3A), thus validating our implementation of the 3’-RACEseq method. For the *AFB2* 5’-cleavage fragment, the 3’-tails also mainly consisted of uridines (Fig EV2A), but we did not detect any clear difference in steady state accumulation of uridylated 5’-cleavage fragments between wild type and *urt1-1* mutants (Fig 3B). In the *heso1-4* mutant background, however, several replicates showed clear accumulation of URT1-dependent, mono-, di- and tri-uridylated species (Fig 3B). Finally, in the *rrp4-2* mutant, we observed a diverse population of oligouridylated fragments (Fig 3C), consistent with their rapid exosome-dependent degradation. To explain these observations, we propose that in wild type, *AFB2* 5’-cleavage fragments are (i) monouridylated by URT1, (ii) extended by HESO1 and probably additional redundantly acting TUTases, and (iii) rapidly degraded by the exosome and, perhaps, additional nucleases such as the RISC-interacting clearing endonucleases (RICE, (Zhang *et al*, 2017)) (Fig 3D). We speculate that partially overlapping functions of the ten TNTases encoded in Arabidopsis may underlie the incompletely penetrant effect of *HESO1* mutation, such that URT1-dependent uridylated species are detectable in only some, but not all, replicates prepared from *heso1-4* mutants. Importantly, preferential monouridylation of *AFB2* 5’-cleavage fragments by URT1 prior to extension by HESO1, but direct uridylation of *MYB33* 5’-cleavage fragments by HESO1 is consistent with established biochemical properties of the two enzymes. URT1 prefers substrates with a 3’-A, and, consequently, catalyses mostly monouridylation (Tu *et al*., 2015b). In contrast, HESO1 prefers substrates with a 3’-U and, consistent with this substrate preference, catalyses formation of longer U-tails (Tu *et al*., 2015b). For *AFB2*, miR393-guided cleavage leaves a 3’-A on the 5’-cleavage fragment, as do miR161.1, miR400, the *TAS2*-derived D9(-) and D11(-) on the *RPF6* 5’-cleavage fragments (Fig 3E and EV2B). In contrast, miR159-guided cleavage of *MYB33* leaves a 3’-U on the 5’-cleavage fragment (Fig 3E). We conclude that our analysis of 3’-end uridylation of *AFB2* 5’-cleavage fragments is consistent with their URT1-dependent uridylation, although this modification has to be understood as one element in a flow of further uridylation and degradation reactions. Taken together with the dependence on the catalytic activity of URT1 of the limitation of miR393-triggered secondary siRNA production, the results strongly indicate that uridylation of the *AFB2* 5’-cleavage fragment limits the secondary siRNA production from the 3’-cleavage fragment, as predicted by the RISC trigger model (Fig 1B).

**Figure 3.**
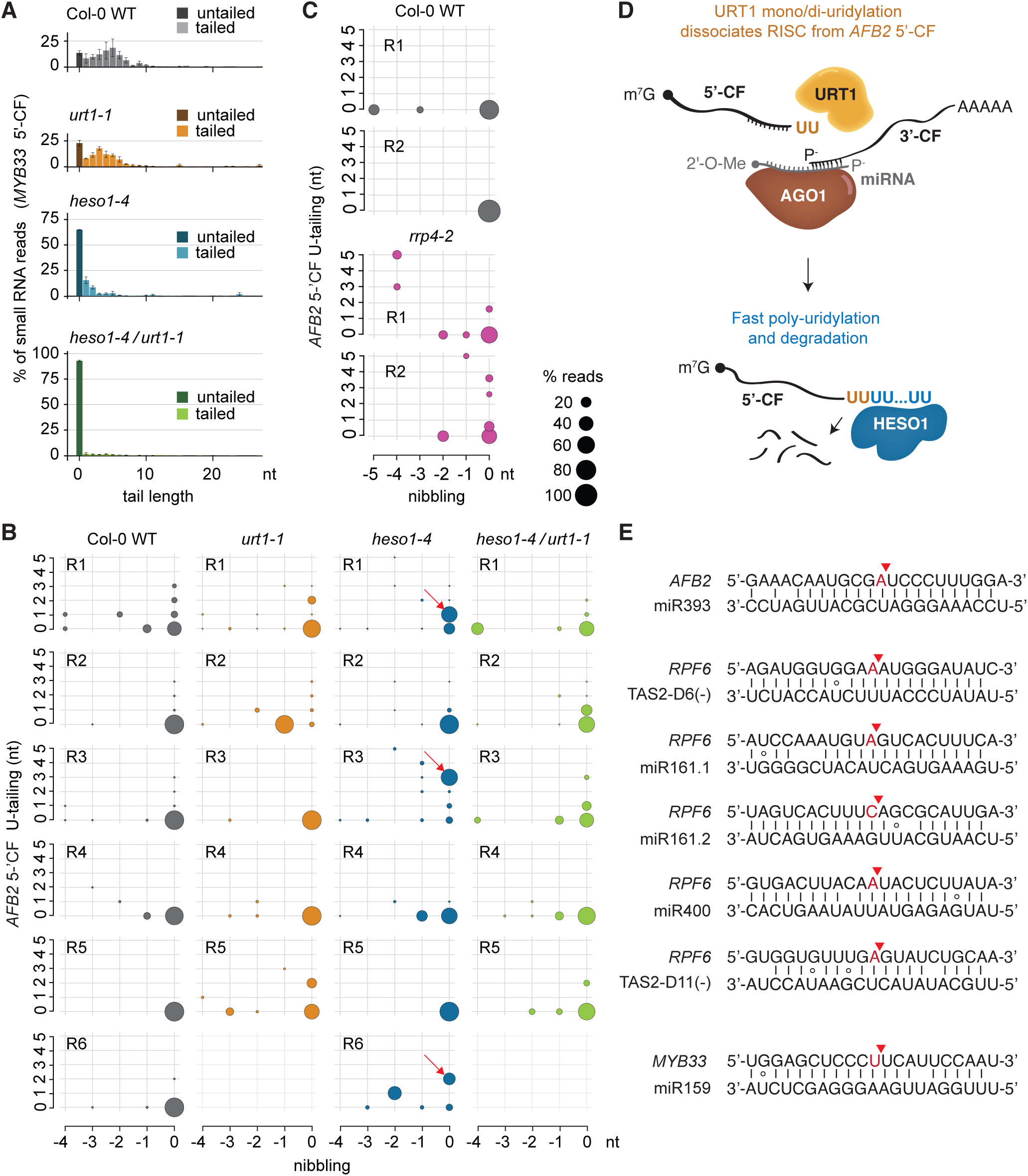
URT1-mediated uridylyl transfer activity on the AFB2 5’ cleavage fragment. **A** Histograms showing the distribution of reads with 0 to 30-nt-long tails on *MYB33* 5’-cleavage fragments (CFs) in the indicated genotypes. Error bars show the standard error of biological triplicates (Col-0 WT and *urt1-1*) or duplicates (*heso1-4* and *heso1-4/ urt1-1*). **B-C** Bubble plots showing the distribution of *AFB2* 5’-CF reads which are up to 4 nt shortened (nibbled) or up to 5 nt longer (tailed). The size of the bubble indicates percentage of nibbled/tailed reads in each genotype and replicate. Only uridine-tails are plotted. **D** Graphical representation of our interpretation of the 3’-RACE-seq results in (B-C). Uridylated *AFB2* 5’-CFs are only observable in *heso1-4* because URT1-mediated mono/di-uridylated intermediates are coupled to subsequent fast uridylation by HESO1 (and probably more TNTases) and degradation by the RNA exosome (and perhaps additional RNA degradation machineries). **E** small RNA target sites within *AFB2, RPF6* and *MYB33* transcripts. Red arrows indicate the cleavage sites. The exposed nucleotides at the 3’ ends of the 5’-CFs are highlighted in red.

### Uncoupling of URT1-mediated uridylation of poly(A) tails and miR393-targeting of AFB2 mRNA

Although the RISC trigger model is consistent with our results on enhanced *AFB2* siRNA production from the 3’-cleavage fragment in *urt1* mutants, we considered an alternative explanation of our results. Since URT1 is also necessary for uridylation of short poly(A) tails transcriptome-wide (Sement *et al*., 2013; Zuber *et al*., 2016), it is possible that the effect on *AFB2* siRNA production mapping to 3’-cleavage fragments is linked to this activity rather than to uridylation of 5’-cleavage fragments. To test this possibility, we analysed *AFB2* secondary siRNA production and uridylation of short poly(A) tails of *AFB2* in the *miR393b* mutant that has >90% reduced levels of miR393 (Si-Ammour *et al*, 2011) (Fig 4A). *AFB2* siRNAs, but not the miR173-triggered, RDR6-dependent siR255 were lost in the *miR393b* mutant (Fig 4A). In contrast, the *AFB2* uridylation pattern on poly(A) tails was similar to wild type. In both the *miR393b* mutant and wild type, those AFB2 poly(A) tails that were further nucleotidylated showed a clear majority of Us at their 3’-ends (Fig 4B) and a similar length distribution of 3’-tails overall (Fig EV3). In contrast, *urt1-1* mutants showed a clear decrease in the presence of Us at the 3’-end of *AFB2* poly(A) tails (Fig 4B), as expected. These results show that uridylation of AFB2 poly(A) tails is not linked to miR393 targeting and that it is not sufficient for *AFB2* siRNA formation, consistent with a scenario in which uridylation of *AFB2* poly(A) tails by URT1 and generation of *AFB2* siRNAs are two separate processes.

**Figure 4.**
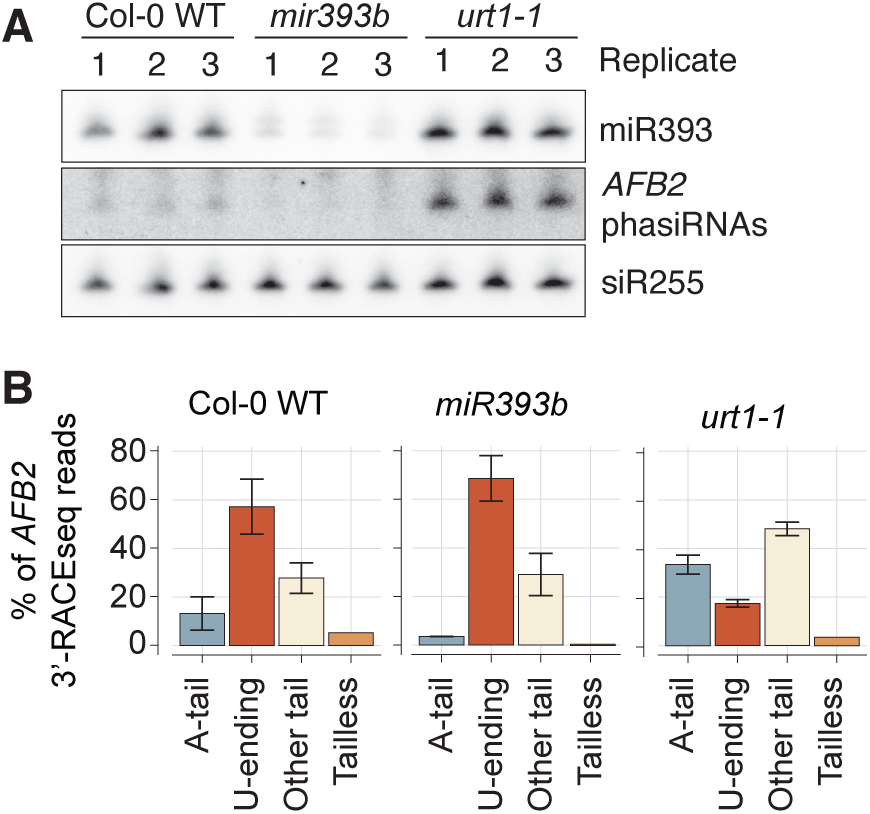
URT1-mediated uridylation of *AFB2* poly(A) tails is independent of miR393-targeting. **A** RNA blot analysis of total RNA from inflorescences of *mir393b* and *urt1-1* mutants in biological triplicates. Radiolabeled probes were hybridized consecutively to the same membrane. siR255 serves as loading control. **B** Bar plots showing the percentage of *AFB2* mRNA tail types in each genotype. The 3’-RACE-seq reads were subcategorized based on their nucleotide content: only consisting of adenosines (A-tail), having a tail of any nucleotide but ending with a uridine (U-ending), all other types of tails (other tail) and reads with no tail (tailless).

### Mutants defective in URT1 and exosome activities show growth phenotypes dependent on DCL2 and RDR6

Since exosome and *urt1* mutants show increased siRNA production from largely distinct subsets of miRNA targets, we next tested whether URT-mediated uridylation of 5’-cleavage fragments and exosomal degradation might act redundantly to limit miRNA-induced secondary siRNA production. We found that both *ski2-5/urt1-1* and *rrp4-2/urt1-1* double mutants showed a clear developmental phenotype that precluded siRNA analysis in flowers, because of growth arrest at the rosette stage (Fig 5A). Interestingly, *ski2-5/heso1-4* and *rrp4-2/heso1-4* double mutants did not exhibit such growth phenotypes (Fig 5A). The developmental phenotype of *rrp4-2/urt1-1* was rescued by transgenic expression of wild type URT1, but not by the catalytically inactive URT1^DADA^ (Fig 5B), demonstrating that loss of URT1 catalytic activity in exosome mutant backgrounds causes growth arrest. We also verified that as in a wild type background, loss of URT1 activity causes increased *AFB2* siRNA populations in the *rrp4-2* mutant background (Fig 5C and Appendix Fig S1).

**Figure 5.**
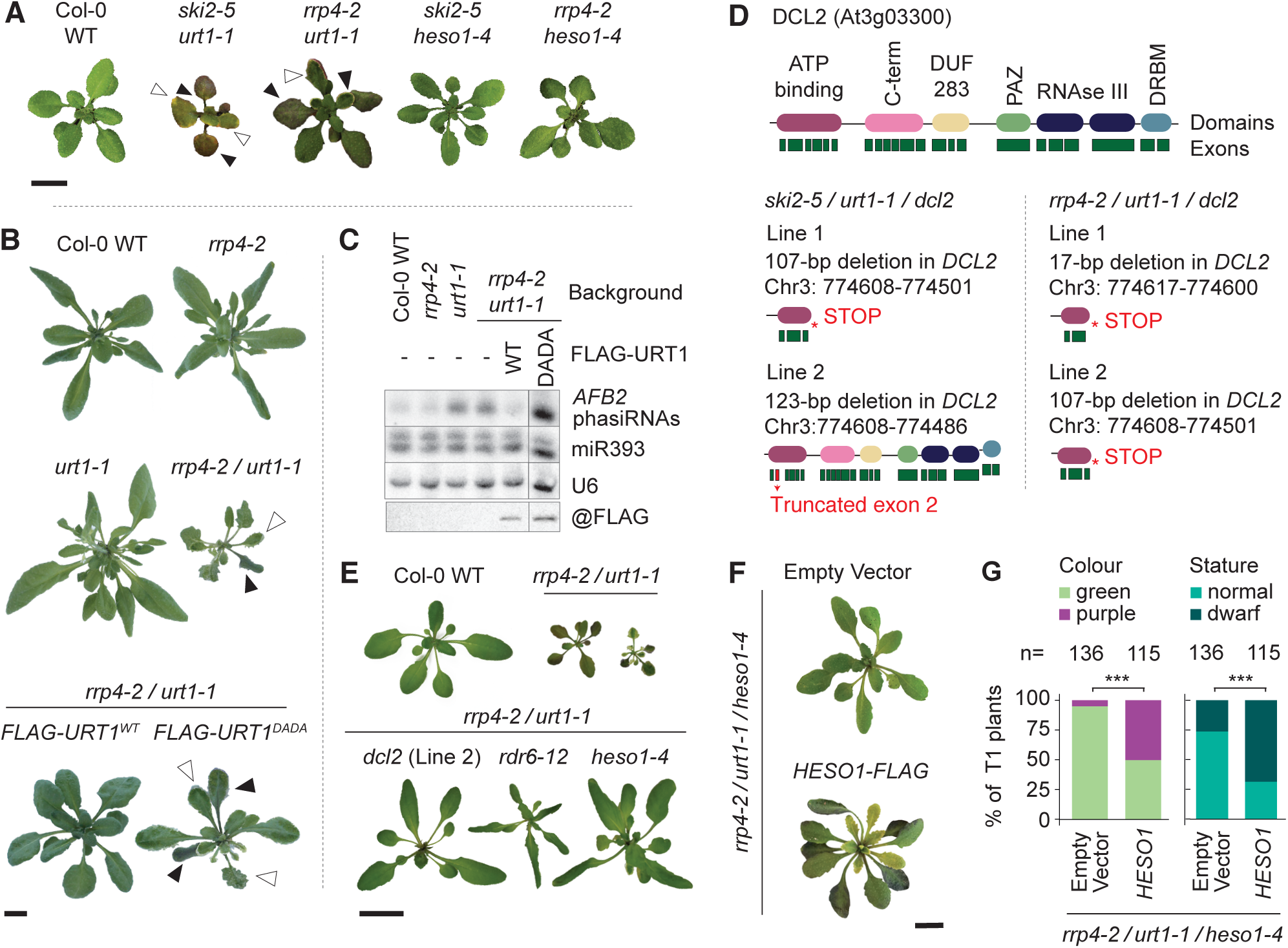
Growth defects in URT1/exosome double mutants are dependent on RDR6, DCL2 and HESO1. **A-B** Rosette phenotypes of the indicated genotypes. White arrows point to leaf serrations, black arrows point to leaves with signs of chlorosis and anthocyanin accumulation. Pictures in A and B were taken 25 and 35 DAG (days after germination) respectively. Scale bars, 1 cm. **C** Top three panels, RNA blot analysis of total RNA from 35-day-old rosettes of the indicated genotypes. Radiolabeled probes were hybridized consecutively to the same blot. U6 serves as loading control. Bottom panel, immunoblot of total protein lysates prepared from rosette tissue. The vertical line indicates the presence of additional lanes removed for presentation purposes. The corresponding full blots are shown in Dataset EV1. **D** Schematic drawing of the *DCL2* gene and encoded protein, and the positions of the truncations caused by the CRISPR-Cas9 deletions engineered here in the indicated backgrounds. **E-F** Rosette phenotypes of the indicated genotypes at 35 (E) and 38 (F) DAG. In E, two individuals with different severity of the *rrp4-2/urt1-1* phenotype are shown. Scale bars, 1 cm. **G** Bar plots showing results of phenotypic analyses of primary transformants (T1s) of *rrp4-2/urt1-1/heso1-4* mutants with either an empty vector or the same vector containing a *HESO1-FLAG* expression construct. Left, percentage of plants showing visible anthocyanin accumulation (purple) in rosette leaves. Right, percentage of plants showing reduced stature (dwarfism). Statistical significance of the difference between Empty Vector and *HESO1-FLAG-*expressing T1 populations was calculated using Student’s *t*-test. *** p < 0.01.

Since growth phenotypes in mutants in RNA metabolism have previously been shown to depend on DCL2 (Bouche *et al*, 2006; Nielsen *et al*, 2024; Zhang *et al*, 2015), we used CRISPR-Cas9 to introduce knockout mutations in *DCL2* in both *ski2-5/urt1-1* and *rrp4-2/urt1-1* mutant backgrounds (Fig 5D). The results showed that knockout of *DCL2* nearly fully suppressed the growth phenotype of *rrp4-2/urt1-1* (Fig 5E), while *ski2-5/urt1-1/dcl2* mutants exhibited partial suppression of the growth phenotype of *ski2-5/urt1-1* (Fig EV4). *rrp4-2/urt1-1/rdr6-12* mutants also showed nearly complete restoration of wild type growth, while retaining the pointy leaf phenotype of *rdr6* mutants (Peragine *et al*, 2004) (Fig 5E) caused by defective tasiRNA biogenesis and overexpression of their transcription factor targets (Adenot *et al*, 2006; Fahlgren *et al*, 2006; Garcia *et al*, 2006; Hunter *et al*, 2006). These results indicate that RDR6-dependent dsRNA production and DCL2-dependent processing are required for the growth phenotypes. Similar characteristics have previously been described for other mutants in basic RNA metabolism, including decapping (*vcs*, phenotypes suppressed in *rdr6* mutants) (Martinez de Alba *et al*, 2015), cytoplasmic 5’-3’ and 3’-5’-exonucleolysis (*xrn4/ski2*, phenotypes suppressed in *rdr6* and *dcl2* mutants) (Zhang *et al*., 2015), and cytoplasmic 5’-3’ exonucleolysis and uridylation (*xrn4/urt1*, phenotypes suppressed in *dcl2/dcl4* double mutants) (Scheer *et al*., 2021). Our results are also in line with the recent report on RDR6-dependent, strong developmental phenotypes in *heso1/urt1/dcl4/ski2* mutants (Wang *et al*, 2022).

### Inactivation of HESO1 suppresses the growth defects of ski2/urt1 and rrp4/urt1 mutants

Because of the partly redundant effects of URT1 and HESO1 in uridylation of miR159-guided *MYB33* 5’-cleavage fragments, we considered the possibility that overlapping effects of the exosome and 3’-uridylation may become clearly visible only upon simultaneous inactivation of both *URT1* and *HESO1* in *ski2* or *rrp4* mutant backgrounds. We therefore constructed triple mutants with the intent to subsequently knock out *DCL2* to recover plants amenable for siRNA analysis in flowers, as described above. Surprisingly, however, inactivation of *DCL2* was not necessary, because the *heso1-4* mutation suppressed the growth phenotype of both *rrp4-2/urt1-1* and *ski2-5/urt1-1* mutants. The suppression of *rrp4-2/urt1-1* phenotypes by *HESO1* inactivation was nearly complete (Fig 5E and Fig EV4), while *ski2-5/urt1-1/heso1-4* exhibited restored growth but a bright yellow color of rosette leaves not seen in *ski2-5/urt1-1* mutants (Fig EV4). When a transgenic copy of *HESO1* was expressed in *rrp4-2/urt1-1/heso1-4*, the developmental defects of *rrp4-2/urt1-1* mutants were restored (Fig 5F,G), confirming that the phenotypic suppression of *rrp4-2/urt1-1* observed in the *heso1-4* mutant background was caused by loss of *HESO1* function. The RDR6-, DCL2-, and HESO1-dependence of the developmental phenotypes of combined exosome/*urt1* mutants will be further treated in subsequent reports. In this paper, we maintain a strict focus on the effect of the exosome and TUTases on miRNA-triggered secondary siRNA production.

### The exosome and HESO1/URT1 redundantly limit secondary siRNA production from a subset of miRNA targets

We next used the *rrp4-2/urt1-1/dcl2* and *rrp4-2/urt1-1/heso1-4* mutants to analyse siRNA populations. Since most secondary siRNAs are DCL4-dependent 21-nt species (Dunoyer *et al*, 2005; Howell *et al*., 2007; Xie *et al*, 2005; Yoshikawa *et al*, 2005), such analyses are still meaningful in the absence of DCL2. We first noticed that simultaneous inactivation of *RRP4* and *URT1* led to ectopic siRNA production from many genes not known to be miRNA targets (Fig EV5), consistent with previous analyses (Li *et al*., 2019; Scheer *et al*., 2021; Vigh *et al*., 2022). A specific focus on miRNA targets showed that simultaneous mutation of *RRP4* and *URT1* resulted in a higher number of miRNA targets with significantly more siRNAs than wild type than the sum of such targets in *rrp4* and *urt1* single mutants (Fig 6A). Gratifyingly, targets with such synergistic effects in *rrp4-2/urt1-1/dcl2* mutants tended to have a 3’-A on their 5’-cleavage fragments (Fig 6B), consistent with URT1-mediated uridylation. Examples of this class include the miR161/miR400 target *RFL9* and the miR399 target *UBC24/PHO2* (Fig 6C). These observations indicate that exosomal degradation and 3’-uridylation indeed exhibit a degree of redundancy in limitation of secondary siRNA production. Nonetheless, most miRNA:target mRNA pairs remain inefficient sources of secondary siRNAs when both exosomal decay and 3’-tailing systems are defective, a point that we treat in detail in the Discussion. Interestingly, for *MYB33* and *TCP4*, whose 5’-cleavage fragments have a 3’-U, clear synergistic effects with *rrp4* were seen only upon inactivation of both *HESO1* and *URT1* (Fig 6C), consistent with the loss of the 3’-U tailing of the *MYB33* 5’-cleavage fragment only upon simultaneous inactivation of *URT1* and *HESO1* (Fig 3A, (Zuber *et al*., 2018)).

**Figure 6.**
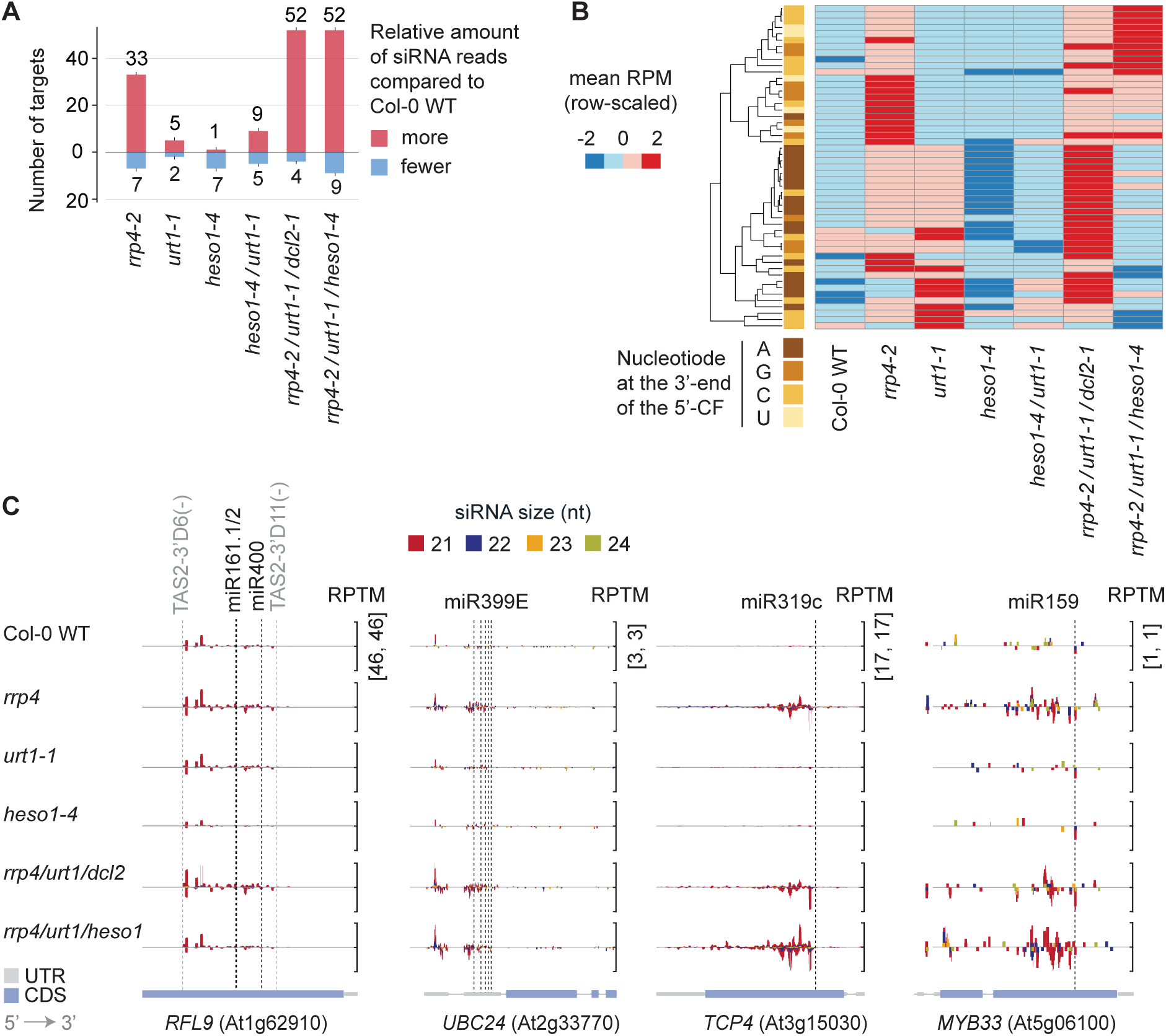
TUTases and the exosome redundantly limit secondary siRNA production from a subset of miRNA targets. **A** Bar plot indicating the number of miRNA targets that give rise to more (red) or less (blue) siRNAs in inflorescences of the indicated genotypes compared to Col-0 WT. **B** Heat map showing row-normalized z-scores of mean RPM (reads per million) of 21-nt siRNAs mapping to miRNA targets in the indicated genotypes, as in Figure 2B. The left column categorizes miRNA targets by the identity of the 3’-nucleotide of the 5’-cleavage fragment. Information on differentially expressed siRNA populations is summarized in Dataset EV1, and sequences of 3’-ends of 5’-cleavage fragments of miRNA targets are listed in Dataset EV2. **C** Examples of miRNA targets in which the accumulation of siRNAs mapping to them is enhanced by simultaneous inactivation of *RRP4* and *URT1* (*RFL9*, *UBC24*), or by simultaneous inactivation of *RRP4*, *URT1* and *HESO1* (*MYB33*, *TCP4*). Representation of mapped reads as in Figure 2C.

### Possible mechanisms of TNTase requirement in secondary siRNA biogenesis

Having established that uridylation of 5’-cleavage fragments limits miRNA-induced secondary siRNA production, in some cases redundantly with the exosome, we moved on to explore possible mechanisms underlying the requirement for HESO1 for normal secondary siRNA production. This requirement was not confined to the *AFB2* and *RPF6* secondary siRNA populations analysed in Fig 2C, but included the abundant miR173-triggered *TAS1B* and *TAS1C* tasiRNAs (Fig 7A). We considered two non-mutually exclusive possibilities. In the first, TNTases would act to extend 3’-ends of single-stranded RNA (ssRNA) templates for RDR6, thereby making them better substrates for the polymerase. Such a mechanism would be reminiscent of the poly(UG) extension of RdRP substrates by the TNTase RDE-3 in *C. elegans* (Shukla *et al*., 2020). In the second, TNTases might act to extend the dsRNA product of RDR6 to make it a better DCL4 substrate. For instance, human Dicer cleaves dsRNA with a variety of 3’-ends, but its specificity and efficiency are optimal with dsRNA substrates with a 2-nt 3’-overhang (Vermeulen *et al*, 2005). Arabidopsis DCL4 also has detectable activity in total lysates against dsRNA with both blunt ends, 1-nt and 2-nt 3’-overhangs (Nagano *et al*, 2014), but the experimental settings employed did not allow estimation of kinetic parameters, leaving open the possibility that processing by DCL4 is also optimal with dsRNA substrates containing 2-nt 3’-overhangs. Arabidopsis RDR6 has weak TNTase activity *in vitro*, but it is unclear whether it can only add one untemplated nucleotide (Curaba & Chen, 2008), as is the case for RDR1 purified from tomato leaves (Schiebel *et al*, 1993b; Schiebel *et al*, 1998). Finally, HESO1 has a modest uridylation activity towards dsRNA *in vitro* that appears to preferentially yield short 1-2 nt tails (Kong *et al*, 2021). This is in contrast to its activity towards ssRNA, where HESO1 efficiently adds long U-tails (Kong *et al*., 2021; Tu *et al*, 2015a; Wang *et al*, 2015). Because of these biochemical properties of DCL4, RDRs, and HESO1, we focused on 2-nt extensions of RDR6 products as a possible mechanism underlying the requirement for HESO1 in secondary siRNA biogenesis using the small RNA-seq data produced in the preceding parts of the study as experimental basis.

**Figure 7.**
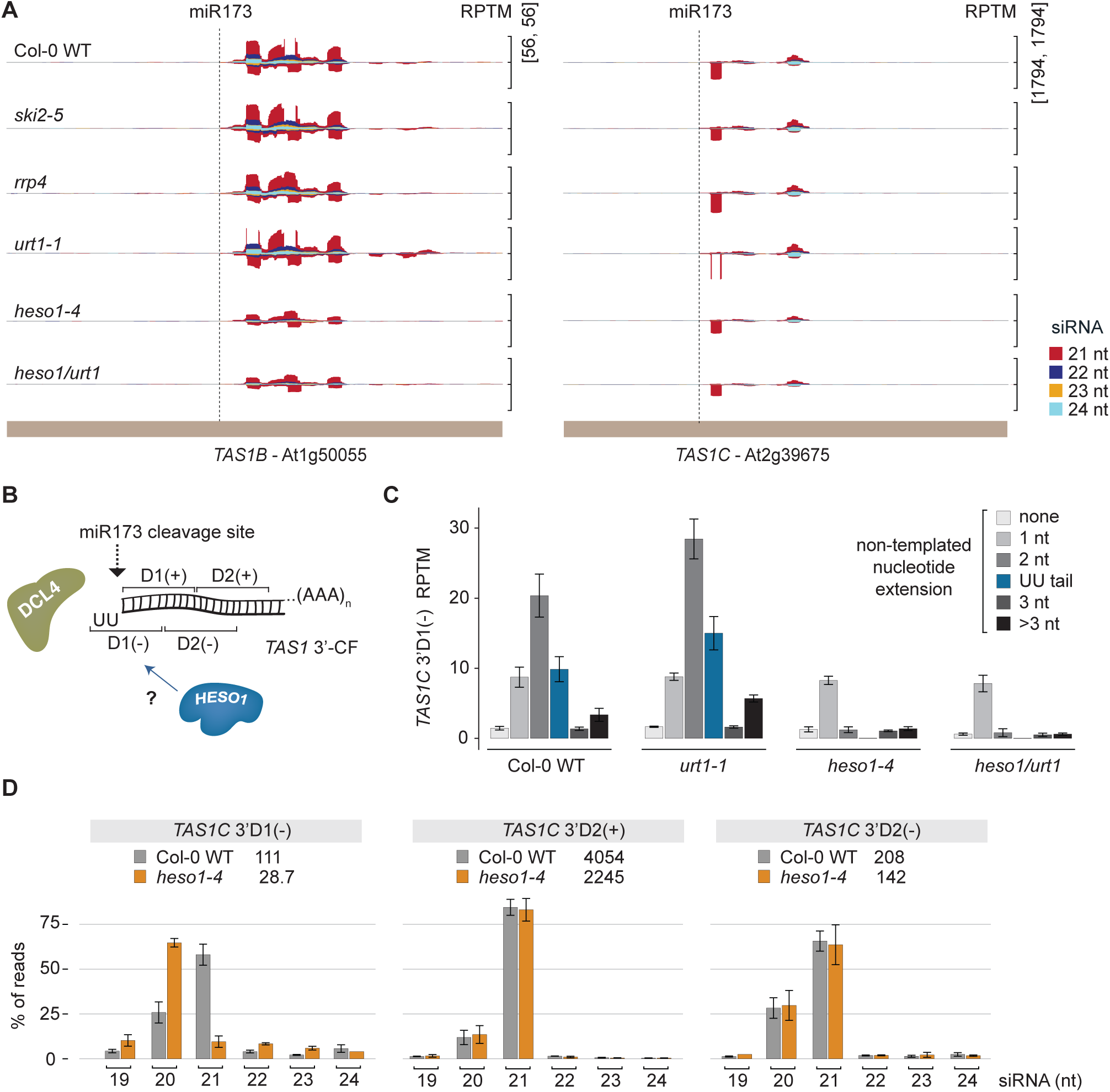
HESO1 is required for 3’-UU extension of RDR6 products. **A** Accumulation of siRNAs derived from *TAS1B* and *TAS1C* represented as in Figures 2C and 6C. **B** Model illustrating a possible role of HESO1 in making 2-nucleotide UU-extensions on RDR6-products to facilitate processing by DCL4. **C** Bar plot showing normalized read counts of the *TAS1C* 3’-D1(-) species in the indicated genotypes. Different *TAS1C* 3’-D1(-) species are differentiated by the presence of untemplated nucleotides at the 3’-end as indicated. RPTM, reads per ten million. **D** Bar plots indicating the size distributions, within the [19-24]-nt range, of three *TAS1C*-derived siRNAs resulting from the first two DCL4 processing events counting from the miR173 cleavage site (*TAS1C* 3’-D1(-) and D2(- and +)) in Col-0 and *heso1-4* mutant libraries. The total number of reads corresponding to the indicated siRNA species is indicated above each bar plot.

### HESO1 is required for 3’-UU extensions of RDR6 dsRNA products

We reasoned that untemplated 3’-extension of RDR6 products should be visible in small RNA-sequencing experiments in the following way: For miRNA targets with 3’-spreading of secondary siRNAs, it should be detectable as a presence of two untemplated 3’-nucleotides on the first antisense siRNA counting from the slicer site, referred to as the 3’-D1(-) siRNA species. Similarly, for miRNA targets with 5’-spreading of secondary siRNAs, it should be detectable as a presence of two untemplated 3’-nucleotides on the first sense siRNA counting from the slicer site, referred to as 5’-D1(+) (Fig 7B). The most abundant miRNA-induced secondary siRNA populations in Arabidopsis are the *TAS1A/B/C* and *TAS2* siRNAs induced by 3’-spreading from the miR173 target site in *TAS1/TAS2* precursor transcripts. In libraries from wild type, only *TAS1C* produced a sufficient number of 3’-D1(-) siRNA reads to complete meaningful analyses (Fig 7C and EV6A). Remarkably, 65% of all *TAS1C* 3’-D1(-) reads had 2-nt extensions in wild type, and of the 16 possible different dinucleotides, nearly 50% were UU (Fig EV6B), consistent with action of a TUTase. Indeed, in the *heso1-4* knockout mutant, the fraction of *TAS1C* 3’-D1(-) reads with 2-nt 3’-extensions was reduced and UU-extensions were completely abolished (Fig 7C), indicating that HESO1 is responsible for the 3’-UU extension of *TAS1C* dsRNA produced by RDR6. Consistent with this difference in 2-nt extensions, the size distributions of the *TAS1C* 3’-D1(-) species were markedly different between wild type and *heso1-4* mutants. In wild type, the predominant size was 21 nt, while it was 20 nt in *heso1-4* (Fig 7D). In addition, the fraction of 19-nt species and of >21-nt species was increased in *heso1-4* (Fig 7D), suggesting reduced specificity of the first processing step. This difference was not caused by generally altered processing by DCL4 in *heso1-4* mutants, because the next siRNA species produced from the cleaved *TAS1C*-derived dsRNA,

*TAS1C* 3’-D2(-) and *TAS1C* 3’-D2(+), did not show any significant difference in size distribution between wild type and *heso1-4* mutants (Fig 7D). We conclude that secondary siRNA biogenesis involves 2-nt extension of the blunt dsRNA produced by RDR6, and that HESO1 contributes a substantial fraction of such extensions, consistent with its requirement for fully efficient production of secondary siRNA from many loci.

## DISCUSSION

Our work defines two distinct steps in secondary siRNA biogenesis in which 3’-uridylation by TUTases plays key roles. The first takes place at the beginning of the process and is important for the decision of whether to recruit RDR6 to an RISC target. This role relies on uridylation of 5’-cleavage fragments produced by miRNA-guided AGO1-catalysed endonucleolysis, seen most clearly here with URT1, although HESO1, and potentially other redundantly acting TNTases, are also likely to be involved. The second takes place at the end of the process, after RDR6 has synthesized dsRNA to be processed by DCL4. This role relies on 2-nt extension of the blunt dsRNA product, involves HESO1, but not URT1, as a predominant player, and is likely to optimize the dsRNA substrate for processing by DCL4. We first discuss the role of the 5’-cleavage fragment uridylation in the context of the overall mechanism of controlled RDR6 recruitment, including the relation between effect sizes of TUTase inactivation observed here and other mechanisms targeting the 5’-cleavage fragment. We then discuss the 3’- end extension of RdRP products discovered here within the broader context of eukaryotic siRNA amplification.

### Limitation of initiation of secondary siRNA formation by uridylation of 5’-cleavage fragments

Our work provides clear examples in which defective uridylation of 5’-cleavage fragments results in enhanced miRNA-triggered secondary siRNA production. We consider these observations to be of substantial importance because they provide evidence for the notion that uridylation of 5’-cleavage fragments limits secondary siRNA production, in some cases redundantly with degradation of 5’-cleavage fragments mediated by the exosome. Nonetheless, the generality of this phenomenon could not be proved through the use of mutants in the exosome and the two best studied TUTases, URT1 and HESO1. We think it plausible that uridylation of 5’-cleavage fragments to limit secondary siRNA production is widespread, but that redundancy in both degradation and uridylation systems underlies the difficulty of detecting this effect, hence explaining our largely sporadic observation of the importance of uridylation of 5’-cleavage fragments for limitation of secondary siRNA production. First, 10 genes encode TNTases in Arabidopsis, of which at least four (NTP2, NTP4, NTP6, NTP7) in addition to HESO1 and URT1 have been shown to have some substrates related to RNA silencing pathways (Kong *et al*., 2021; Song *et al*, 2019). Thus, for many miRNA targets, 3’-tailing of 5’-cleavage fragments may simply be catalysed by other nucleotidyl transferases in the absence of URT1 and HESO1. Second, at least two classes of nucleases in addition to the exosome, the RICE endonucleases and the 3’-5’ exonuclease ATRM2 act on RISC-associated RNAs (Wang *et al*, 2018; Zhang *et al*., 2017), raising the possibility that the combined action of these enzymes is sufficient to limit secondary siRNA production from many miRNA targets, even with the reduced exosome and TUTase activities obtained here through inactivation of *URT1*, *HESO1*, *SKI2* and *RRP4*. Third, we analysed the effects of the genetic knockouts on secondary siRNA production using total inflorescence tissue containing a mix of many different cell types. It is possible that this relatively crude experimental design precludes observation of more widespread effects if they are only apparent in cell types with particularly high expression of the genes in question. In this case, a simple dilution effect of the number of siRNA reads detected would compromise the ability to detect statistically significant differential siRNA expression at the depth of sequencing used here.

### Uridylation of 5’-cleavage fragments and the key importance of RISC dissociation from target RNA

The results of this study are consistent with predictions of the RISC trigger model for secondary siRNA production, because they provide examples of uridylation of 5’-cleavage fragments that limits secondary siRNA production from 3’-cleavage fragments. This behavior is coherent with the RISC trigger model if one assumes that uridylation of 5’-cleavage fragments accelerates RISC dissociation from the RISC:target RNA complex post cleavage, as outlined in Figure 1. We note that the idea of accelerated RISC dissociation from target RNAs by uridylation of 5’-cleavage fragments also offers an explanation for uridylation of 5’-cleavage fragments produced by siRNA-guided cleavage in metazoans (Shen & Goodman, 2004) that do not encode RdRPs. Since RISC dissociation from target RNA may be rate-limiting for endonucleolysis in a multiturnover setting (Deerberg *et al*, 2013), the uridylation of 5’-cleavage fragments may simply ensure sufficient RISC activity in these species. Such a fundamental implication of uridylation of 5’-cleavage fragments by TUTases associated with endonucleolytic RISCs would offer an attractive explanation for the necessity to evolve protective 3’-end methylation of all plant miRNAs and siRNAs, metazoan transposon-targeting Piwi-associated RNAs (piRNAs) and *Drosophila* Ago2-associated siRNAs, but not metazoan miRNAs (Horwich *et al*, 2007; Kamminga *et al*, 2010; Kirino & Mourelatos, 2007; Li *et al*, 2005; Pastore *et al*, 2021; Saito *et al*, 2007; Yu *et al*, 2005).

### Requirement for 3’-nucleotidylation in secondary siRNA production

Our results also provide several prominent examples of facilitation of miRNA-triggered secondary siRNA production by HESO1. The discovery that HESO1 mediates 3’-UU extension of RDR6-produced dsRNA derived from *TAS1C*, and that other dinucleotides may also be added, presumably by other TNTases, prompts the following interpretation: TNTases, in particular HESO1, constitute a core element in secondary siRNA formation by preparing an optimal DCL4 substrate with a 2-nt 3’-overhang from the dsRNA produced by RdRP. Consistent with this interpretation, the RdRP-containing complex (RDRC) in the yeast *S. pombe* contains the TNTase Cid12 in addition to the polymerase itself (Motamedi *et al*., 2004). It is now an attractive possibility that the biochemical basis for the absolute requirement of Cid12 for siRNA amplification in *S. pombe* (Motamedi *et al*., 2004) is to prepare appropriate Dcr1 substrates coupled to dsRNA synthesis by the RdRP. Interestingly, plant RdRP-containing complexes may also contain TNTases, because early purifications of RDR1 from tomato leaves took considerable care to separate it from a co-fractionating TNTase activity (Schiebel *et al*, 1993a). In addition, HESO1-mediated mono- or diuridylation of RNA polymerase IV/RDR2 products has recently been described, although it is unclear whether this 3’-end extension influences processing by the cognate DICER-LIKE ribonuclease, DCL3 (Ren *et al*, 2023).

The action of HESO1 and other TNTases at the level of dsRNA may not be the full explanation for their requirement for secondary siRNA biogenesis, however. The single-stranded RDR6 template is also an attractive target of TNTase action, as 3’-extension might promote RDR6-mediated conversion into dsRNA. A number of different observations suggests that TNTase-catalysed 3’-extension of single-stranded RdRP template RNAs may indeed be involved in siRNA production. First, in *C. elegans*, (UG)_n_-tailing by the mixed-nucleotide TNTase RDE-3 is required for conversion of single-stranded RNAs into dsRNA by RdRPs (Shukla *et al*., 2020). Second, also in *C. elegans*, full siRNA amplification depends on the endonuclease RDE-8, and RDE-8-mediated cleavage is required for target fragment uridylation and siRNA amplification by RdRPs (Tsai *et al*, 2015). Third, RDR6 activity is inhibited by a consecutive stretch of 8 adenosines (Baeg *et al*, 2017), excluding the possibility that polyadenylated 3’-cleavage fragments serve as direct substrates of RDR6. Thus, 3’-modification may be involved in channeling substrates to RDR6, making analysis of 3’-ends of RDR6-substrates in combinations of mutants with inactivation of RDR6 and TNTases a promising future line of study to fully elucidate the requirement of TNTases for secondary siRNA biogenesis in plants.

## MATERIALS AND METHODS

### Plant material, growth conditions and genotyping

All plants used in this study are of the ecotype Columbia (Col-0). T-DNA insertion mutants in *SKI2* (AT3G49690: *ski2-2* (SALK_129982), *ski2-5* (SALK_118529)), URT1 (AT2G45620: *urt1-1* (SALK_087647C) and *urt1-2* (WISCDSLOXHS208_08D)) and *MIR393B* (AT:*mir393b* (SALK_018966) were previously described (Branscheid *et al*., 2015; Sement *et al*., 2013; Si-Ammour *et al*., 2011), as were the point mutant alleles *rrp4-2/sop2* (Hematy *et al*., 2016) and *rdr6-12* (Peragine *et al*., 2004). Mutants in *HESO1* (AT2G39740: *heso1-3* (SALK_044238C) and *heso1-4* (GK_369H06)) were identified in this study from the SALK and GABI-KAT collections (Alonso *et al*, 2003; Rosso *et al*, 2003). The T-DNA insertion mutants and the *rdr6-12* mutant were all obtained from the Nottingham Arabidopsis Stock Centre. The *rrp4-2* mutant was a gift from Kian Hématy.

Molecular analyses were carried out with inflorescence tissue with exception of the sRNA northern blots in Figures 2E and 5C, which were carried out in rosettes. All seeds were sterilized and germinated on plates containing 1x Murashige & Skoog medium (#M0222.0050, Saveen og Werner ApS, Denmark), supplemented with 1% (w/v) sucrose and 0.8% (w/v) agar. Seedlings were transferred to soil (Plugg/Såjord [seedcompost]; SW Horto, Bramming, Denmark) 10 days after germination (DAG). In experiments where *ski2/urt1* double and *ski2/urt1/dcl2* triple mutants were included, all seedlings were germinated directly on soil. After transfer to soil, plants were grown in a Percival growth chamber for an additional 4 weeks under the following conditions: 16 h light (Master TL-D 36W/840 and 18W/840 bulbs (Philips); 130 mmol m^−2^ s^−1^, 21°C, 60% relative humidity)/ 8 h darkness (16°C, 60% relative humidity). Inflorescences and rosettes for northern blots, western blots and sequencing libraries were collected and snap-frozen in liquid nitrogen.

Each T-DNA insertion mutant was genotyped with two PCR reactions: a PCR reaction with a set of primers flanking the insertion site for wild type allele detection and a PCR reaction with an outward left border primer and one of the two flanking primers for T-DNA allele detection. Genotyping of *rdr6-12* and *rrp4-2* was done as described (Arribas-Hernandez *et al*., 2016; Vigh *et al*., 2022). All primers used for genotyping in this study were purchased from TAG Copenhagen A/S and their sequences are listed in Data Set EV3.

### Double and triple mutant construction

The *heso1-4/urt1-1* double mutant was constructed by crossing *urt1-1* to *heso1-4.* F1 plants were self-pollinated and in the F2 population, double mutants were identified by genotyping. *urt1-1* was also crossed to *rrp4* and *ski2-5,* but here, all double mutants in the F2 population died at the rosette stage. Therefore, seeds from *urt1/rrp4(+/-)* and *urt1/ski2(+/-)* plants were kept as the final seed stocks and double mutants were identified by genotyping in the beginning of each experimental setup. The triple mutant *rrp4/urt1-1/heso1-4* was constructed by crossing *heso1-4* to *urt1/rrp4(+/-).* F1 plants were self-pollinated and in the F2 population, triple homozygous mutants were identified by genotyping. *heso1-4* was also crossed to *urt1/ski2(+/-),* but here, all triple mutants in the F2 population turned completely yellow and died at the rosette stage. Therefore, seeds from *heso1-4/urt1-1/ski2-5(+/-)* plants were kept as the final seed stock. The triple mutant *rrp4/urt1-1/rdr6-12* was constructed by crossing *rdr6-12* to *urt1/rrp4(+/-).* F1 plants were self-pollinated and in the F2 population, triple homozygous mutants were identified by genotyping. Genotyping primers are listed in Dataset EV3.

### CRISPR-Cas9 DCL2 cloning

*DCL2* (AT3G03300) was knocked out using the CRISPR-Cas9 vector-system described in (Tsutsui & Higashiyama, 2017). Three different guide oligos were inserted in each their pKAMA-ITACHI Red (pKIR) vector, a vector encoding RFP-Cas9, the following way: First, the pKIR vector was digested with AaRI (Thermo Scientific^TM^), dephosphorylated and gel-purified. Meanwhile, a top and a bottom strand of the guide oligos were synthesized separately and next, hybridized and phosphorylated using PNK and PNK buffer A (Thermo Scientific^TM^). The digested plasmid was used in T4 (Thermo Scientific^TM^) DNA ligation reactions with hybridized and phosphorylated guide oligos. pKIR vectors with correct guide insertions were transformed into *A. tumefaciens* cells.

Simultaneous transformation of *urt1/rrp4(+/-)* and *urt1/ski2(+/-)* plants with a mix of three different *Agrobacterium* cultures each carrying one of the three plasmids for Cas9-guide RNA expression was performed with the floral dip method (Clough & Bent, 1998). Primary transformants (T1s) were screened for deletions in *DCL2* with a PCR reaction spanning the DNA regions wherein the guide RNAs direct Cas9 dsDNA breakage. The PCR fragments with deletions were gel-purified and sent for Sanger sequencing with the forward primer from the PCR reaction. Seeds of T1 plants with sequenced deletions were sorted using a fluorescence stereomicroscope to select those in which the seed coat fluorescence marker linked to Cas9 had been lost by segregation. T2 plants were genotyped for deletions in both *DCL2* alleles and for *rrp4* or *ski2-5* homozygosity. Two different transgenic lines (L1 and L2) of each original transformation were kept. T2 seeds of *rrp4/urt1/dcl2* L1 and L2 were plated on MS-agar plates with hygromycin to ensure loss of the Cas9-encoding transgene. Afterwards, the lines were backcrossed once to minimize CRISPR-Cas9 off-targets in the transgenic lines. All top and bottom guide oligos and the genotyping primers are listed in Dataset EV3.

### Cloning of FLAG-URT1^WT^, FLAG-URT1^DADA^, and HESO1^WT^-FLAG

To construct *proURT1::FLAG::URT1^WT^::terURT1* and *proHESO1::HESO1^WT^-FLAG::terHESO1* transgenes, two PCR fragments compatible for downstream USER cloning (Bitinaite & Nichols, 2009) were amplified from Col-0 WT genomic DNA with the KAPA HiFi Hotstart Ready-Mix from KAPA Biosystems. The transgenes were split into two PCR fragments either at the N- or C terminus depending on the desired position of the FLAG-tag in the final construct. The FLAG-tag was encoded in the primers. For the transgene constructions, we used the pCAMBIA3300U vector adapted to USER cloning reactions (Nour-Eldin *et al*, 2006), a kind gift from Barbara A. Halkier. The transgene encoding catalytically URT1 (*proURT1::FLAG::URT1^DADA^::terURT1*) was also constructed by USER cloning by splitting one of the two PCR reactions described above to create USER fragments with flanking regions at the mutation site. The D491A and D493A mutations were encoded in the PCR primers. All USER primers are listed in Dataset EV3.

### Plant transformation and selection of transgenic lines

Transgenic plant lines were constructed with the floral dipping method (Clough & Bent, 1998). Flowering plants of either *urt1-1, urt1-2, urt1/rrp4(+/-)*, *urt1/ski2(+/-)* or *rrp4/urt1/heso1* genotypes were dipped in a suspension of *Agrobacterium tumefaciens* GV3101 bacteria in 5% sucrose and 0.05% Silwet L-77 with an OD_600_ ∼ 0.8-1.2. The seeds of the dipped plants were spread directly on soil and sprayed with glufosinate ammonium (0.2 g/L) on day 7, 10 and 14. Around 20-25 T1 plants were kept for screening for single T-DNA insertion in the T2 and T3 generations and for protein expression levels. 2-3 independent transgenic lines of the T3 generation with similar expression levels were kept.

### RNA extraction

Total RNA from inflorescences was extracted with TRI Reagent (#T9424, Sigma-Aldrich Denmark A/S) according to the manufacturer’s instructions. The RNA was dissolved in either 50% formamide for northern blots or in sterilized water for small RNA libraries and RT-PCR analysis.

### sRNA northern blot

10-20 μg of total RNA was suspended in sRNA loading buffer (20 mM HEPES pH7.8, 1 mM EDTA, 50 % formamide, 3 % glycerol, bromophenol blue (BPB) and xylene cyanol). The RNA was denatured 65°C for 10 min. prior to loading in a pre-warmed 18% acrylamide:bis 19:1, 7% urea, 0.5 x TBE gel. Electrophoresis in 0.5 TBE at 100 V was performed until the BPB reached the bottom of the gel. The gel was blotted onto an Amersham Hybond-NX membrane at 80 V in cold 0.5 TBE for an hour. The membrane was chemically crosslinked with EDC (1-ethyl-3-(3-dimethylaminopropyl) carbodiimide) for 1-2 hours at 60 °C. After crosslinking, the membrane was rinsed with water and incubated at 42 °C for 20 min. in 8 mL PerfectHyb^TM^ Plus Hybridization buffer. After 20 minutes, a radiolabelled probe was added directly in the buffer and the membrane was incubated overnight (O/N) with the probe at 42 °C. The following day, the membrane was washed with 2xSSC (0.3 M NaCl, 30 mM sodium citrate, 2 % SDS) three times of minimum 10 min. at 42 °C. The washed membrane was developed with phosphorimaging. If more than one probe was hybridized, the membrane was stripped with 0.1% boiling SDS in between hybridizations. Detection of a single miRNA or siRNA species was done by end-labelling complementary oligonucleotides, by incubation with 2.5 µCi γ-^32^P-ATP (PerkinElmer) and 1U T4 polynucleotide kinase (ThermoScientific) according to the manufacturer’s instructions. Labelled probes were separated from unincorporated γ^32^P-ATP by gel filtration using G-25 MicroSpin Columns (GE Healthcare Life Sciences). The U6 probe was end-labelled the same way.

The probe for detection of *AFB2* secondary siRNAs was made by incorporation of α^32^P-dCTP in a Prime-a-Gene reaction from Promega. The reaction was carried out as described in the manual from the manufacturer only with substitution of dCTP from the kit with α^32^P-dCTP. The template for the Prime-a-Gene reaction was a ∼ 300 nt long dsDNA product obtained from a PCR reaction from plant cDNA with *AFB2* specific primers. 50 ng of the dsDNA template was used in the Prime-a-Gene reaction. Probes were cleaned from unincorporated α^32^P-dCTP with Sephadex G-50 columns. All sequences of oligonucleotides for miRNA/siRNA detection and the primers used to obtain the *AFB2* secondary siRNA probe are listed in Dataset EV3.

### FLAG-HRP western blot

For screening of protein levels in the transgenic *FLAG-URT1*, *FLAG-URT1^DADA^*, and *HESO1-FLAG* lines, total lysates were made in the following way: Flower or seedling tissue was ground in liquid nitrogen using a mortar. 400 μL of cold IP buffer (50 mM Tris-HCl pH 7.5, 150 mM NaCl, 10% glycerol, 5 mM MgCl_2_ and 0.1% Nonidet P40) was supplemented with protease inhibitor (Roche Complete) and added to 100 mg of ground tissue. All samples were vortexed until homogenous. The lysates were cleared by centrifugation at 16,000 *g* for 15 minutes at 4°C. The cleared lysates were transferred to new Eppendorf tubes and mixed with Laemmli sample buffer (70 mM Tris-HCl pH 6.8, 10% glycerol, 1% LDS and 0.01 % BPB). The samples were heated at 85°C for 5 minutes prior to SDS-PAGE. The proteins in the SDS-PAGE gel were transferred onto a nitrocellulose membrane (Amersham Protran Premium) for 45 minutes at 80V in the cold. The membrane was afterwards blocked in 5% skimmed milk in PBS-T (1 x PBS, 0.05 % Tween-20) for 30 minutes. The blocked membrane was incubated over night with monoclonal Anti-FLAG M2 antibody conjugated to horse radish peroxidase (HRP) (Sigma) in 5 % milk. The following day, the membrane was washed with PBS-T 5 x 5 minutes and the HRP-signals were developed with enhanced chemiluminescence (ECL). If signals were weak, SuperSignal^TM^ West Femto Maximum Sensitivity Substrate from ThermoFisher was used.

FLAG-Immunopurification (IP) was performed prior to western blotting (Figure 1E) in the following way: 100 mg of ground seedling tissue was homogenized in 1 mL IP-buffer and cleared lysate prepared as described above were transferred to a pre-cooled Eppendorf tube and incubated with Anti-FLAG M2 affinity resin (Sigma). 10 μL of bead-slurry was incubated with 1 mL of plant lysate. The incubation was performed at 4°C for 1-2 hours under slow rotation. The beads were washed in wash buffer (50 mM Tris-HCl pH 7.5, 500 mM NaCl, 10% glycerol, 5 mM MgCl_2_ and 0.1% Nonidet P40) 3 x 5 minutes before Laemmli sample buffer was added to the washed beads. The beads were boiled in Laemmli buffer for 5 minutes at 85°C prior to SDS-PAGE.

### sRNA sequencing library organization and construction

Sequencing libraries were constructed from three biological replicates of each genotype. Each biological replicate consisted of inflorescences collected from four individual plants grown in the same pot. Plants of all genotypes were grown in parallel. The resulting data from Col-0, *ski2-5* and *rrp4-2* mutants have been reported (Vigh *et al*., 2022).

1 μg of total inflorescence RNA was used as input in each library. Libraries were generated using the NEBNext Multiplex Small RNA Library Prep Set (#E7300S, New England Biolabs) according to the manufacturer’s instructions. The indexed cDNA libraries were size selected on a 6% polyacrylamide gel as described in the NEBNext protocol. All size-selected libraries were analysed using an Agilent Bioanalyser, quantified with Qubit measurements (#Q32854, Invitrogen) and single-end sequenced on an Illumina Nextseq 500 with SE75_HI chemistry (#FC-404-2005, Illumina). 1% of spike-in PhiX Control v3 (#15017666, Illumina) was also loaded.

### Adapter trimming and mapping conditions

Adapter sequence trimming and sRNA read mapping were done as described (Vigh et al., 2022).

### Differential expression analysis

The pipeline used for the DESeq2 analysis has been described (Vigh *et al*., 2022). The code for producing Figures 2 and 6 can be found on the GitHub repertoire https://github.com/MariaLouisaVigh/HESO1URT1 together with a list of all genes with differentially expressed siRNA levels in each mutant (Dataset EV1).

### RACEseq library construction

The genotypes and replicates used in the RACEseq experiments were Col-0 WT (6 replicates), *urt1-1* (5 replicates), *heso1-4* (6 replicates), *heso1/urt1* (5 replicates). In an individual RACEseq experiment, libraries from Col-0 WT (2 replicates) and *rrp4-2* (2 replicates) were constructed. Each biological replicate was derived from inflorescences of four plants grown in the same pot. The RNA was extracted with TRI Reagent (#T9424, Sigma-Aldrich Denmark A/S), resuspended in water followed by a polysaccharide precipitation as described (Arribas-Hernandez *et al*., 2016). Finally, the RNA was once again resuspended and purified with phenol:chloroform:isoamyl alcohol. RNA integrity was assessed on a Bioanalyser RNA Nano Chip (Agilent Technologies). Twenty pmol of a 3’-ssDNA adapter (custom order, IDT), previously described in (Zuber *et al*., 2018) was ligated to 10 μg of total RNA from each sample listed above. When incubating total RNA and the adapter with 20 U T4 DNA ligase (ThermoScientific), the adapter ligates to all RNA with a hydroxyl 3’-OH end due to an adenyl pyrophosphoryl moiety on the adapter 5’-end. To separate excess adapter from successfully ligated RNA, samples were purified using NucleoSpin® RNA Clean-up columns (Macherey Nagel). Next, 2 x 20 μl reactions of 2 μg of denatured RNA incubated with 50 pmole of 3’-RT oligo, 10 mM dNTP mix, 200 U of SuperScript IV reverse transcriptase (Invitrogen^TM^), 1X SuperScript RT buffer, 0.1 M DTT and 40 U Ribolock (ThermoScientific) were incubated at 50°C for 10 minutes. The reverse transcriptase was then inactivated at 80°C for 10 min. The cDNA was purified with phenol:chloroform:isoamyl alcohol and the aqueous phase was EtOH-precipitated. The cDNA was resuspended in 8μl of water.

Two rounds of nested PCR reactions were then performed. The first PCR contained 2 μl of cDNA, 10 pmol of a gene-specific primer (*MYB33*_5’CF/*AFB2*_5’CF/*AFB2*_3’CF), 10 pmol of RACEseq_rev1 primer, 12.5 nmol dNTP mix, 1U of Phusion DNA Polymerase and 1 x Phusion HF buffer (ThermoScientific). The PCR specifications were 94 °C 30 sec, (94°C 20 sec, 50°C 20 sec, 72°C 30 sec) x 20 cycles, 72 °C 30 sec. Thereafter, 4 x 20 μl reactions for each gene-specific PCR reaction were set up using 1 μl of PCR reaction 1, 10 pmole of a new gene-specific primer, 10 pmole of TRUseq RPI primer, 12.5 nmole of dNTP mix, 1 U Phusion DNA Polymerase and 1 x Phusion HF buffer (ThermoScientific). The PCR specifications were 94°C 1 min, (94°C 30 sec, 56°C 20 sec, 72°C 30 sec) x 25 cycles, 72 °C 30 sec. The four identical PCR reactions were pooled, and the libraries were PCR purified using Beckman Coulter^TM^ Agencourt AMPure XP beads following the manual from the provider. The libraries were assessed with Agilent Bioanalyser on a DNA 1000 chip and the concentration was determined with Qubit HS DNA chemistry (Invitrogen^TM^). Libraries were pooled in equimolar ratios and sequenced on a NextSeq (v3 chemistry) MidOutput flow cell with 41 x 111 bp settings. 50% of phiX was added to compensate for the poor read diversity of poly(A)-tails. Sequences of all adapters and primers used for the library construction are listed in Dataset EV3.

### RACEseq pipeline

Fastq files were sorted based on their gene-specific primer used in the 2^nd^ PCR reaction, which is for *MYB33*: 5’-AAGAATTCTCGTCGCCTGAA-3’, *AFB2*_5’CF: 5’-GGTCTCCTTACAGACCAAGT-3’ and *AFB2*_3’CF: 5’-GATTCTTCTGTGAAAGCCATTCTG-3’. Next, only read 2 fastq files beginning with the 3’-adapter delimiter sequence followed by 15 random bases (Unique Molecular Identifier, UMI) were kept and were deduplicated using the UMI. Thereafter, the 3’-end of the deduplicated reads were trimmed using *Cutadapt v.2.4* (Martin, 2011) for adapter leftovers (TTCAGGCGACGAGAATTCTT). The final output is a .txt file wherein each line represents a tail of a unique read. To obtain such a file, two rounds of *Cutadapt* with special command lines were used to first trim away the gene body until the CS (in the case of *AFB2* 3’-CF the gene body was cleaved until the end of the cDNA sequence given in TAIR). Unspecific reads included due to off-targets of the nested gene-specific primers were removed in the same trimming step by the command *-- discard-untrimmed.* In the second round of *Cutadapt* trimming, *AFB2*_5’CF and *MYB33*_5’CF reads that have not been cleaved were removed by searching for reads with a sequence running over the CS followed by the command *-- discard-trimmed.* For the nibbling and tailing analysis, the same pipeline was used but the last Cutadapt steps were changed so that they trimmed the gene body until -10 nt upstream of the CS. Lastly, the .txt files were imported in R and deduplicated once again with a custom R script that only allows UMI that differ from each other in minimum 13 of the 15 random bases. The pipeline together with a R script of the downstream analyses of the .txt files can be found on Github: https://github.com/MariaLouisaVigh/HESO1URT1

### Analysis of 3’-end nucleotides of 5’-cleavage fragments

A FASTA file of 326 mature miRNA sequences was downloaded from *miRbase* (Griffiths-Jones, 2004). The reverse complement base of the 11^th^ nt was written in a separate column with the use of the R package *Biostrings v2.58.0* (https://bioconductor.org/packages/Biostrings). The column with the last nt of the 5’-CF was joined to the in-house miRNA target list. The final list is summarized in Dataset EV2.

### Non-templated sRNA tail analysis

In Figure 7, 3’-D1-(-), 3’-D2-(-) and 3’-D2-(+) reads from *TAS1A/B/C* and *AFB2* were retrieved by filtering fastq files for reads with a 19-nt match in sequence to the small RNA species in question. Whether a read was trimmed or tailed and which nucleotides were added was computed in R.

## DATA AVAILABILITY

Small RNA-seq and 3’-RACE-seq data have been deposited in the Gene Expression Omnibus under accession number GSE267947 and are publicly available as of May 22, 2024. All code used to analyse sequencing data is available in the Github repository: https://github.com/MariaLouisaVigh/HESO1URT1

## ACKNOWLEDGMENTS

We thank Dominique Gagliardi for *urt1* mutant seeds and for advice on 3’-RACE-seq analysis. Freja Asmussen and August Sveen Jespersen are thanked for technical assistance, and Theo Bölsterli and René Hvidberg Petersen and their teams are thanked for plant care.

## AUTHOR CONTRIBUTIONS

MLV conducted all experiments and data analysis included in this manuscript, AT discovered the increased siRNA populations of miR393 targets in *urt1* mutants by analysis of a separate small RNA-seq data set prepared by LA-H, LA-H and PB conceived the initial project objective of investigating defective limitation of secondary siRNA production in TUTase mutants, PB supervised the project and wrote the paper with input from all authors.

---

**Appendix Figure S1.**
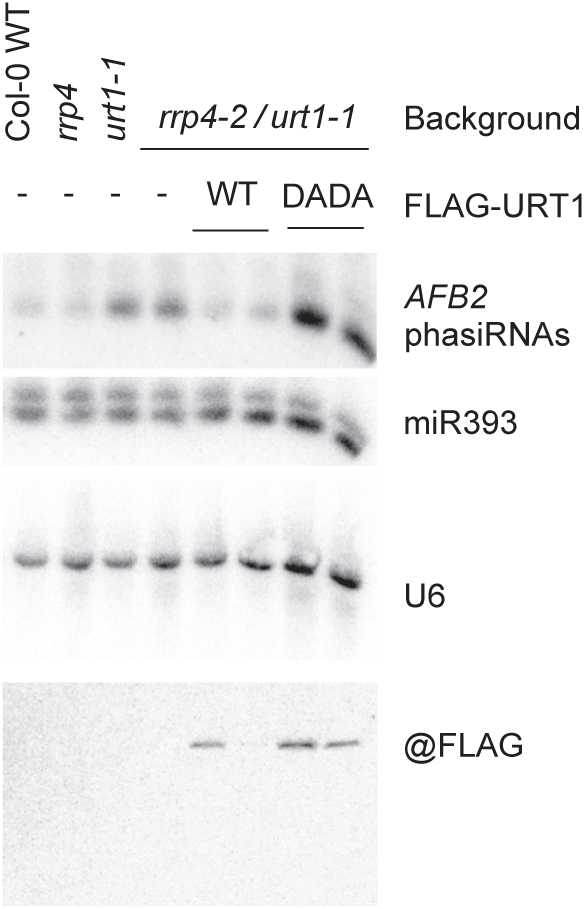
URT1 uridylyl activity is required for limiting the production of *AFB2*-derived secondary siRNAs in *rrp4/urt1 double mutants* (Uncropped version of Figure 5C) Top three panels, RNA blot analysis of 10 μg total RNA extracted from 35-day-old rosettes of the indicated genotypes. Radiolabeled probes were hybridized consecutively to the same blot. U6 serves as loading control. Bottom panel, protein blot of total lysates of rosette tissue. Two independent transgenic lines are shown for *urt1-1*/*FLAG-URT1^WT^* and *urt1-1*/*FLAG-URT1^DADA^*.

**Figure EV1.**
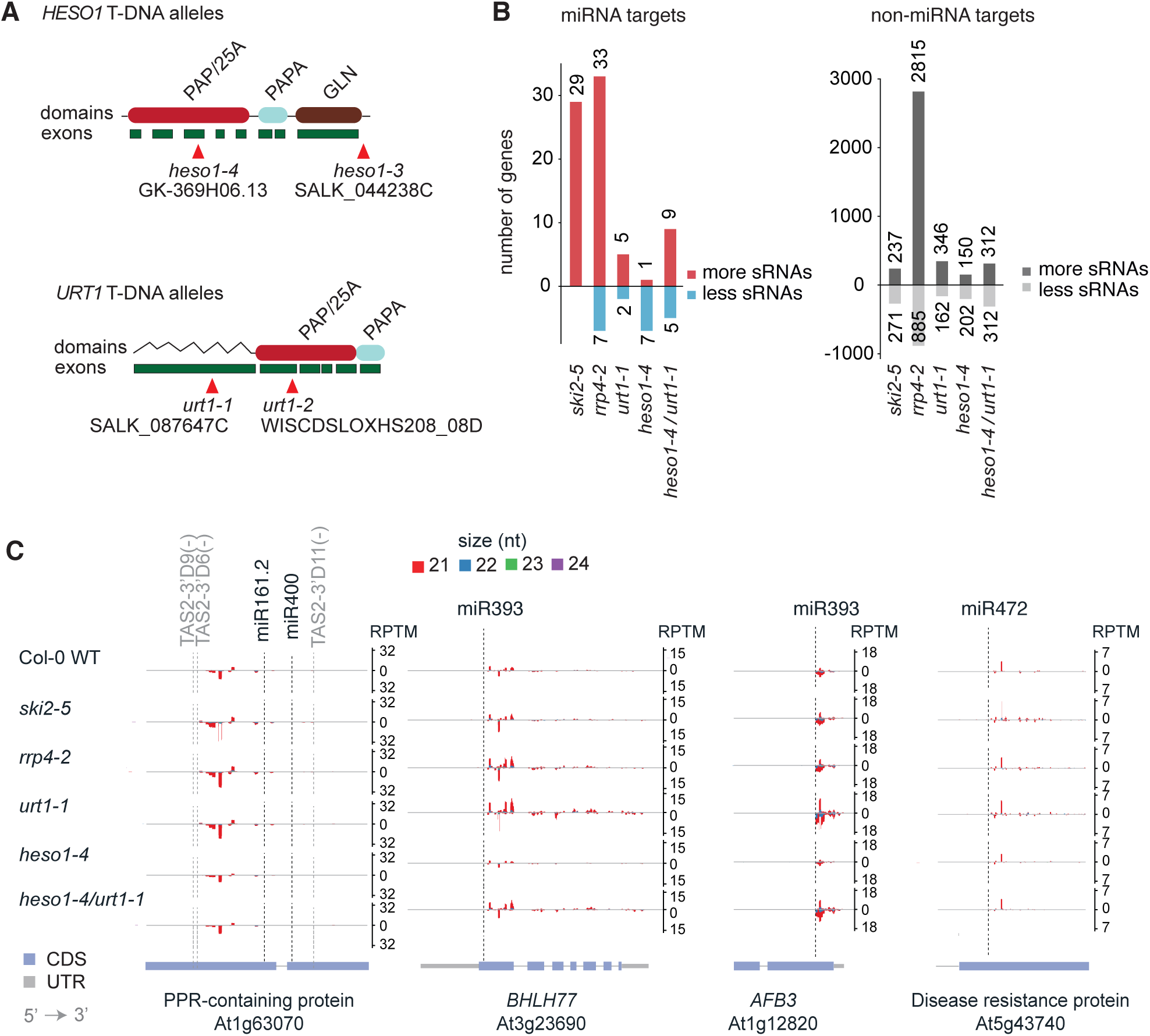
siRNA production from miRNA targets in *urt1* and *heso1* mutants (Supports Figure 2) **A** Schematic illustration of the *HESO1* and *URT1* genes, the encoded proteins, and the positions of the T-DNA insertions used for loss-of-function analysis in this study. PAP, poly(A) polymerase domain, PAPA, PAP-Associated domain, GLN, glutamine-rich region. The wavy line in URT1 indicates an intrinsically disordered region. **B** Bar plots indicating the number of loci with more or less siRNAs in inflorescences of the indicated geno-types compared to wild type. **C** Examples of accumulation of 21- to 24-nt siRNAs mapping to miR393, miR472, and miR161/miR400/-*TAS1*-tasi targets in the indicated genotypes. RPTM, reads per ten million. Reads above the X axes are in sense orientation, and below are antisense. Dashed lines indicate small RNA cleavage sites. Transcript architecture is represented below.

**Figure EV2.**
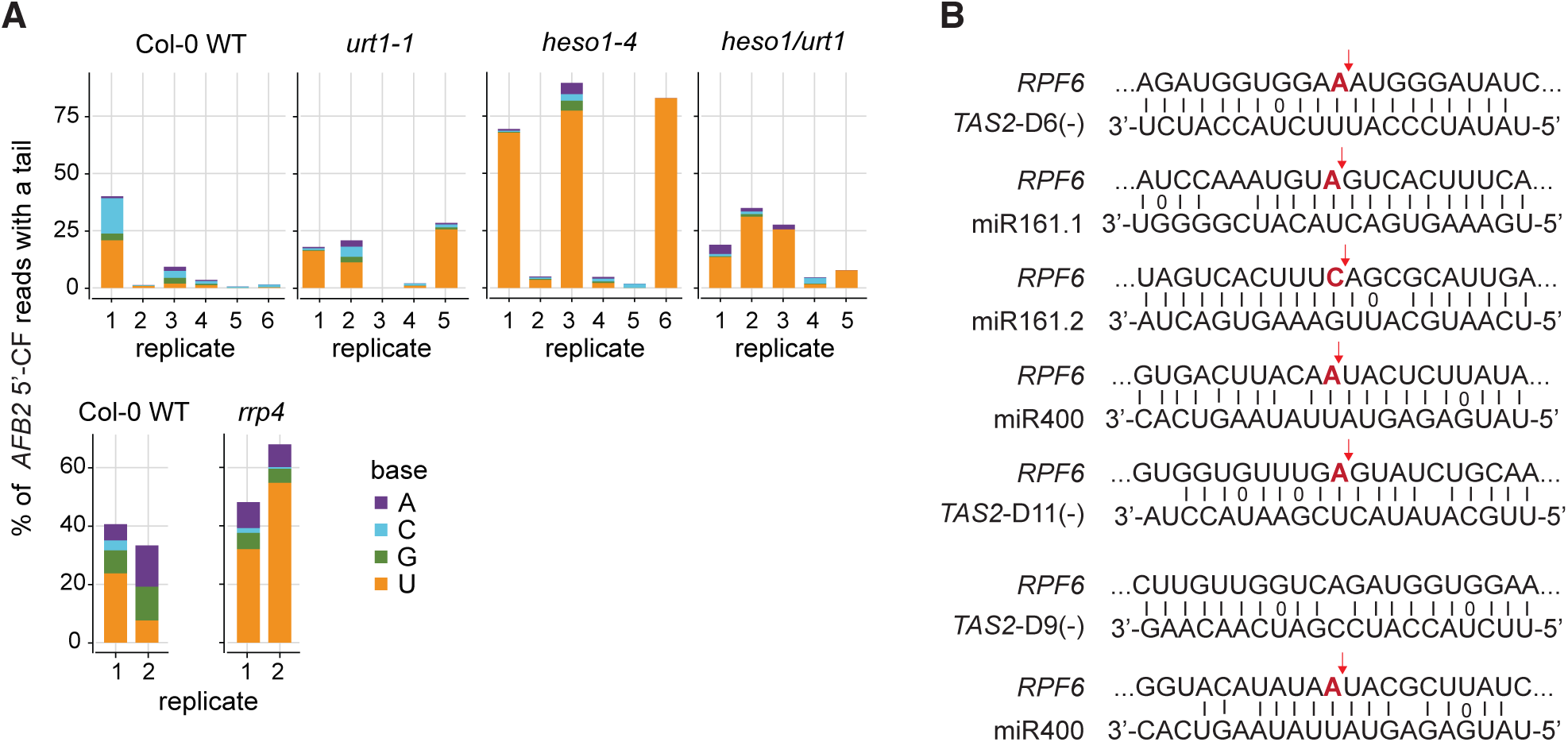
Nucleotide composition of non-templated tails on the 3’ end of the *AFB2* 5’-cleavage fragment, and predicted target sites in *RPF6* mRNA (Supports Figure 3) A Bar plots depicting the percentage of tailed *AFB2* 5’-cleavage fragment (CF) reads obtained by 3’-RACE-seq with a tail and the relative content of adenosine (A), cytidine (C), guanosine (G) and uridine (U) in these tails. All biological sample replicates in two independent experiments, one with Col-0 WT, *urt1-1*, *heso1-4* and *heso1-4/urt1-1* (top), and the other with Col-0 WT and *rrp4-2* (bottom), are plotted individually. **B** All predicted target sites in *RPF6* ranked by their Expectation value (1.5 - 5) from psRNAtarget (Dai & Zhao, 2011). All *RFP6* target site sequences are represented in 5’-to-3’ orientation. Dai X, Zhao PX (2011) psRNATarget: a plant small RNA target analysis server. Nucleic acids research 39: W155-W159

**Figure EV3.**
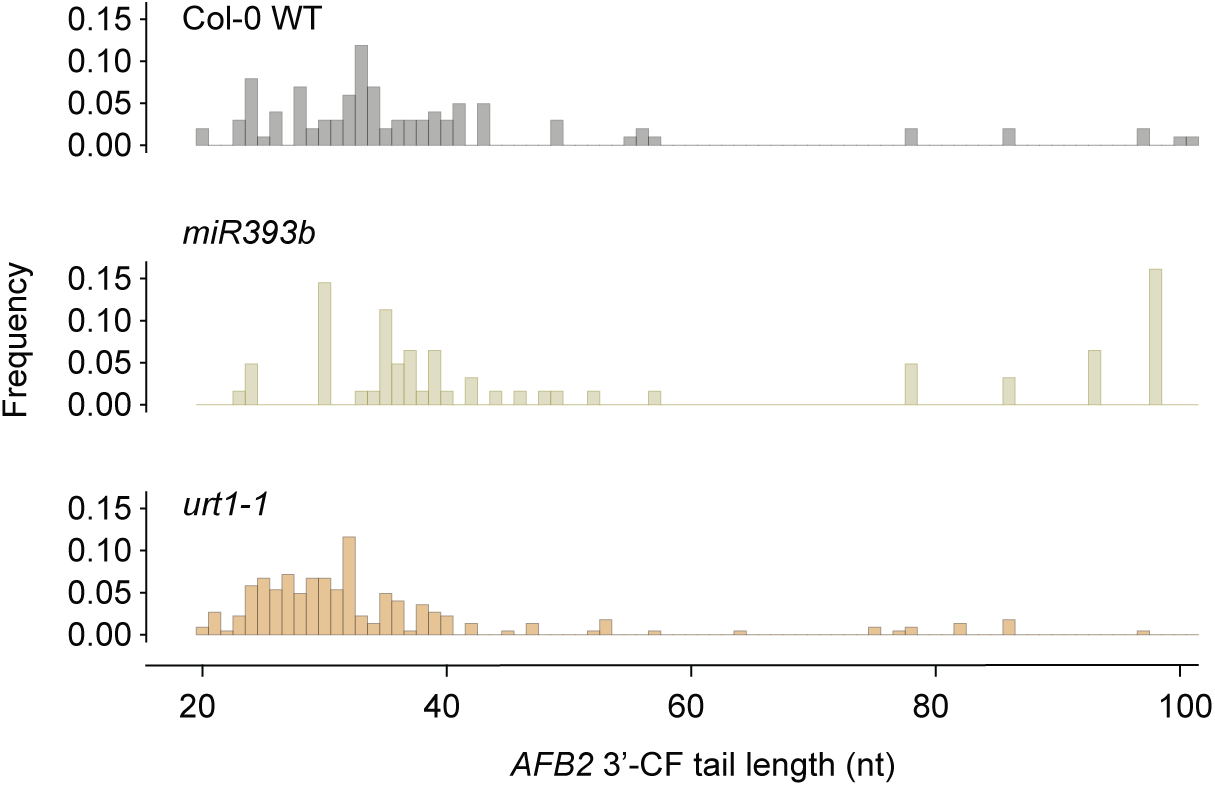
Distribution of *AFB2* poly(A) tail length (Supports Figure 4) Histogram depicting the distribution of poly(A) tail lengths, between 0-120 nt, on the *AFB2* 3’-CF in Col-0 WT, *urt1-1* and *mir393b* mutants. Reads from biological triplicates are merged.

**Figure EV4.**
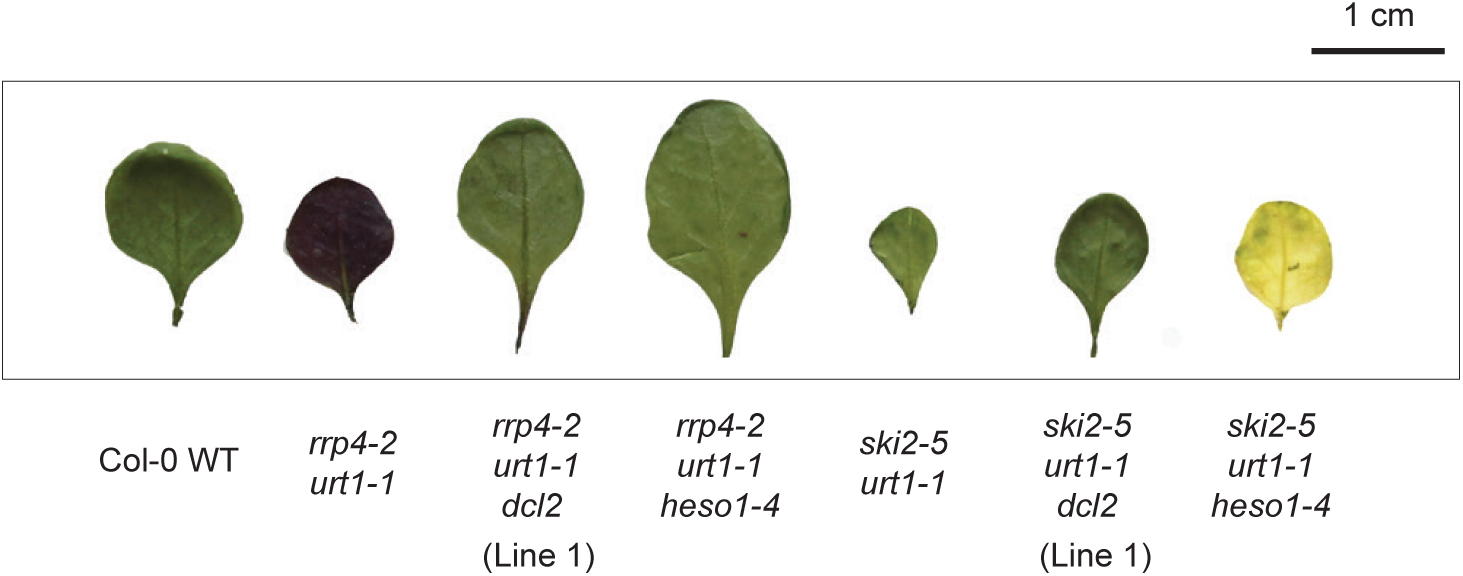
comparison of phenotypes in triple mutants (Supports Figure 5) Photographs of the abaxial side of one of the first true leaves of 25-day-old plants of the indicated genotypes.

**Figure EV5.**
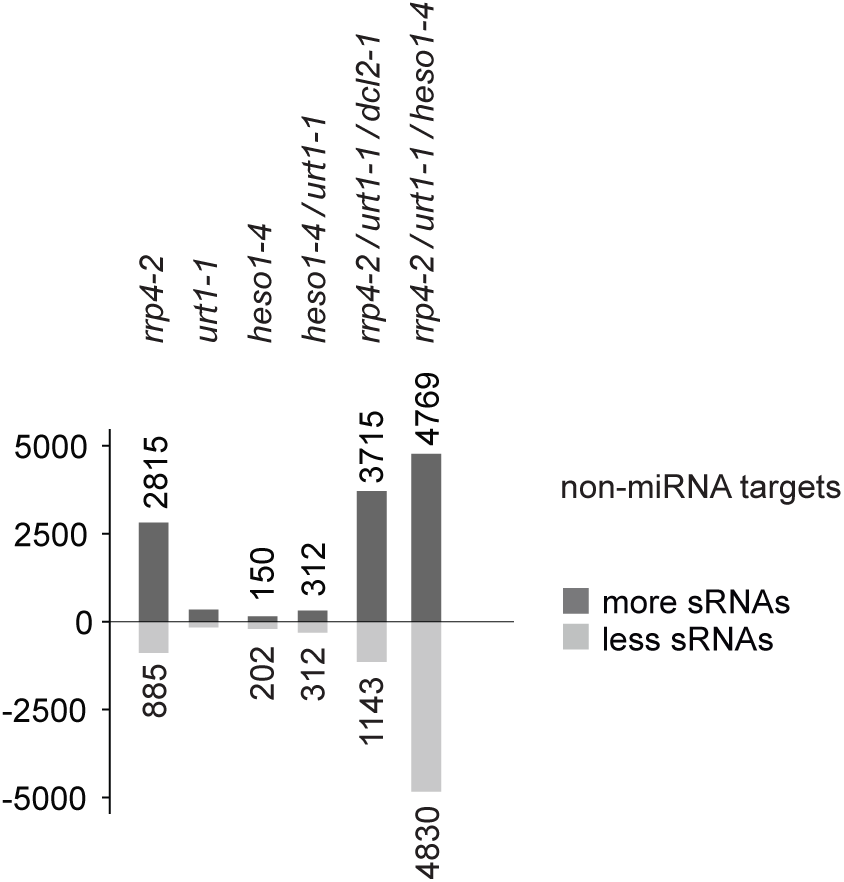
siRNA production from non-miRNA targets in triple mutants (Supports Figure 6) Bar plot showing number of non-miRNA targets with more (dark grey) or less (light grey) siRNAs in inflorescences of the plants of the indicated genotypes compared to Col-0 wild type individuals.

**Figure EV6.**
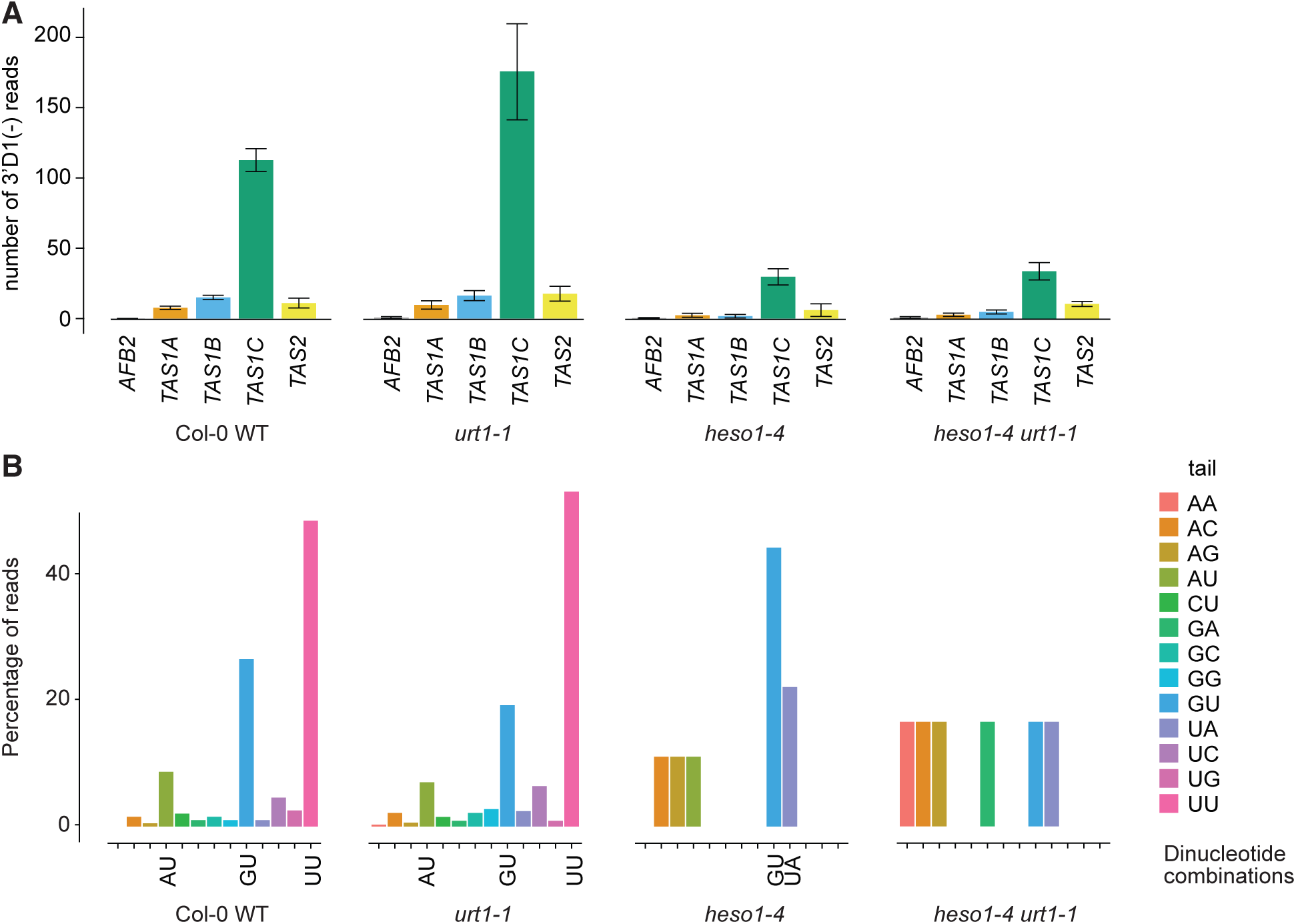
Abundance of 3’D1(-) secondary siRNA species (Supports Figure 7) **A** Bar plot showing number of 3’D1(-) reads mapping to *AFB2, TAS1A/B/C* and *TAS2* transcripts in libraries from the four indicated genotypes. Error bars indicate the standard error of read counts in biological triplicates. **B** Percentage distribution of dinucleotide combinations of 3’-tails on *TAS1C* 3’D1(-) reads. CA, CC and CG dinucleotides were not detected in any of the libraries from the four indicated genotypes. The bars indicate the fraction of each dinucleotide extension summed over all reads across the three biological triplicates.

## REFERENCES

Adenot X, Elmayan T, Lauressergues D, Boutet S, Bouche N, Gasciolli V, Vaucheret H (2006) DRB4-dependent TAS3 trans-acting siRNAs control leaf morphology through AGO7. Current biology : CB 16: 927–932

Allen E, Xie Z, Gustafson AM, Carrington JC (2005) microRNA-directed phasing during trans-acting siRNA biogenesis in plants. Cell 121: 207–221

Alonso JM, Stepanova AN, Leisse TJ, Kim CJ, Chen H, Shinn P, Stevenson DK, Zimmerman J, Barajas P, Cheuk R et al (2003) Genome-wide insertional mutagenesis of Arabidopsis thaliana. Science 301: 653–657

Arribas-Hernandez L, Marchais A, Poulsen C, Haase B, Hauptmann J, Benes V, Meister G, Brodersen P (2016) The Slicer Activity of ARGONAUTE1 Is Required Specifically for the Phasing, Not Production, of Trans-Acting Short Interfering RNAs in Arabidopsis. The Plant cell 28: 1563–1580

Axtell MJ, Jan C, Rajagopalan R, Bartel DP (2006) A two-hit trigger for siRNA biogenesis in plants. Cell 127: 565–577

Axtell MJ, Westholm JO, Lai EC (2011) Vive la différence: biogenesis and evolution of microRNAs in plants and animals. Genome biology 12: 221

Baeg K, Iwakawa H-o, Tomari Y (2017) The poly(A) tail blocks RDR6 from converting self mRNAs into substrates for gene silencing. Nature Plants 3: 17036

Bartel DP (2004) MicroRNAs: genomics, biogenesis, mechanism, and function. Cell 116: 281–297

Bernstein E, Caudy AA, Hammond SM, Hannon GJ (2001) Role for a bidentate ribonuclease in the initiation step of RNA interference. Nature 409: 363–366

Bitinaite J, Nichols NM (2009) DNA cloning and engineering by uracil excision. Current protocols in molecular biology / edited by Frederick M Ausubel *[et al]* Chapter 3: Unit 3 21

Boccara M, Sarazin A, Thiébeauld O, Jay F, Voinnet O, Navarro L, Colot V (2014) The Arabidopsis miR472-RDR6 Silencing Pathway Modulates PAMP- and Effector-Triggered Immunity through the Post-transcriptional Control of Disease Resistance Genes. PLOS Pathogens 10: e1003883

Bouche N, Lauressergues D, Gasciolli V, Vaucheret H (2006) An antagonistic function for Arabidopsis DCL2 in development and a new function for DCL4 in generating viral siRNAs. EMBO Journal 25: 3347–3356

Branscheid A, Marchais A, Schott G, Lange H, Gagliardi D, Andersen SU, Voinnet O, Brodersen P (2015) SKI2 mediates degradation of RISC 5’-cleavage fragments and prevents secondary siRNA production from miRNA targets in Arabidopsis. Nucleic acids research 43: 10975–10988

Cerutti H, Casas-Mollano JA (2006) On the origin and functions of RNA-mediated silencing: from protists to man. Current Genetics 50: 81–99

Chen H-M, Chen L-T, Patel K, Li Y-H, Baulcombe DC, Wu S-H (2010) 22-nucleotide RNAs trigger secondary siRNA biogenesis in plants. Proceedings of the National Academy of Sciences 107: 15269–15274

Clough SJ, Bent AF (1998) Floral dip: a simplified method for Agrobacterium-mediated transformation of Arabidopsis thaliana. Plant J 16: 735–743

Cuperus JT, Carbonell A, Fahlgren N, Garcia-Ruiz H, Burke RT, Takeda A, Sullivan CM, Gilbert SD, Montgomery TA, Carrington JC (2010) Unique functionality of 22-nt miRNAs in triggering RDR6-dependent siRNA biogenesis from target transcripts in Arabidopsis. Nature structural & molecular biology 17: 997–1003

Curaba J, Chen X (2008) Biochemical Activities of *Arabidopsis* RNA-dependent RNA Polymerase 6. Journal of Biological Chemistry 283: 3059–3066

Dalmay T, Hamilton A, Rudd S, Angell S, Baulcombe DC (2000) An RNA-dependent RNA polymerase gene in Arabidopsis is required for posttranscriptional gene silencing mediated by a transgene but not by a virus. Cell 101: 543–553

De Almeida C, Scheer H, Zuber H, Gagliardi D (2018) RNA uridylation: a key posttranscriptional modification shaping the coding and noncoding transcriptome. Wiley Interdiscip Rev RNA 9

Deerberg A, Willkomm S, Restle T (2013) Minimal mechanistic model of siRNA-dependent target RNA slicing by recombinant human Argonaute 2 protein. Proceedings of the National Academy of Sciences 110: 17850–17855

Deleris A, Gallego-Bartolome J, Bao J, Kasschau KD, Carrington JC, Voinnet O (2006) Hierarchical action and inhibition of plant Dicer-like proteins in antiviral defense. Science 313: 68–71

Dunoyer P, Himber C, Voinnet O (2005) DICER-LIKE 4 is required for RNA interference and produces the 21-nucleotide small interfering RNA component of the plant cell-to-cell silencing signal. Nature genetics 37: 1356–1360

Fagard M, Boutet S, Morel J-B, Bellini C, Vaucheret H (2000) AGO1, QDE-2, and RDE-1 are related proteins required for post-transcriptional gene silencing in plants, quelling in fungi, and RNA interference in animals. Proc Natl Acad Sci USA 97: 11650–11654

Fahlgren N, Montgomery TA, Howell MD, Allen E, Dvorak SK, Alexander AL, Carrington JC (2006) Regulation of AUXIN RESPONSE FACTOR3 by TAS3 ta-siRNA Affects Developmental Timing and Patterning in Arabidopsis. Curr Biol 16: 939–944

Garcia D, Collier SA, Byrne ME, Martienssen RA (2006) Specification of leaf polarity in Arabidopsis via the trans-acting siRNA pathway. Current biology : CB 16: 933–938

Griffiths-Jones S (2004) The microRNA Registry. Nucleic acids research 32: D109–D111

Hematy K, Bellec Y, Podicheti R, Bouteiller N, Anne P, Morineau C, Haslam RP, Beaudoin F, Napier JA, Mockaitis K et al (2016) The Zinc-Finger Protein SOP1 Is Required for a Subset of the Nuclear Exosome Functions in Arabidopsis. PLoS genetics 12: e1005817

Heo I, Ha M, Lim J, Yoon M-J, Park J-E, Kwon SC, Chang H, Kim VN (2012) Mono-Uridylation of Pre-MicroRNA as a Key Step in the Biogenesis of Group II let-7 MicroRNAs. Cell 151: 521–532

Heo I, Joo C, Kim Y-K, Ha M, Yoon M-J, Cho J, Yeom K-H, Han J, Kim VN (2009) TUT4 in Concert with Lin28 Suppresses MicroRNA Biogenesis through Pre-MicroRNA Uridylation. Cell 138: 696–708

Hernandez-Pinzon I, Yelina NE, Schwach F, Studholme DJ, Baulcombe D, Dalmay T (2007) SDE5, the putative homologue of a human mRNA export factor, is required for transgene silencing and accumulation of trans-acting endogenous siRNA. Plant J 50: 140–148

Horwich MD, Li C, Matranga C, Vagin V, Farley G, Wang P, Zamore PD (2007) The Drosophila RNA Methyltransferase, DmHen1, Modifies Germline piRNAs and Single-Stranded siRNAs in RISC. Curr Biol 17: 1265–1272

Howell MD, Fahlgren N, Chapman EJ, Cumbie JS, Sullivan CM, Givan SA, Kasschau KD, Carrington JC (2007) Genome-wide analysis of the RNA-DEPENDENT RNA POLYMERASE6/DICER-LIKE4 pathway in Arabidopsis reveals dependency on miRNA- and tasiRNA-directed targeting. The Plant cell 19: 926–942

Hunter C, Willmann MR, Wu G, Yoshikawa M, de la Luz Gutierrez-Nava M, Poethig SR (2006) Trans-acting siRNA-mediated repression of ETTIN and ARF4 regulates heteroblasty in Arabidopsis. Development 133: 2973–2981

Ibrahim F, Rohr J, Jeong WJ, Hesson J, Cerutti H (2006) Untemplated oligoadenylation promotes degradation of RISC-cleaved transcripts. Science 314: 1893

Iwakawa H-o, Lam AYW, Mine A, Fujita T, Kiyokawa K, Yoshikawa M, Takeda A, Iwasaki S, Tomari Y (2021) Ribosome stalling caused by the Argonaute-microRNA-SGS3 complex regulates the production of secondary siRNAs in plants. Cell reports 35

Jauvion V, Elmayan T, Vaucheret H (2010) The Conserved RNA Trafficking Proteins HPR1 and TEX1 Are Involved in the Production of Endogenous and Exogenous Small Interfering RNA in Arabidopsis The Plant cell 22: 2697–2709

Kamminga LM, Luteijn MJ, den Broeder MJ, Redl S, Kaaij LJT, Roovers EF, Ladurner P, Berezikov E, Ketting RF (2010) Hen1 is required for oocyte development and piRNA stability in zebrafish. The EMBO journal 29: 3688–3700

Kirino Y, Mourelatos Z (2007) Mouse Piwi-interacting RNAs are 2′-O-methylated at their 3′ termini. Nature structural & molecular biology 14: 347–348

Kong W, Dong X, Ren Y, Wang Y, Xu X, Mo B, Yu Y, Wang X (2021) NTP4 modulates miRNA accumulation via asymmetric modification of miRNA/miRNA* duplex. Science China Life Sciences 64: 832–835

Li J, Yang Z, Yu B, Liu J, Chen X (2005) Methylation protects miRNAs and siRNAs from a 3’-end uridylation activity in Arabidopsis. Current biology : CB 15: 1501–1507

Li T, Natran A, Chen Y, Vercruysse J, Wang K, Gonzalez N, Dubois M, Inzé D (2019) A genetics screen highlights emerging roles for CPL3, RST1 and URT1 in RNA metabolism and silencing. Nature Plants 5: 539–550

Lisitskaya L, Aravin AA, Kulbachinskiy A (2018) DNA interference and beyond: structure and functions of prokaryotic Argonaute proteins. Nature communications 9: 5165

Liu J, Carmell MA, Rivas FV, Marsden CG, Thomson JM, Song JJ, Hammond SM, Joshua-Tor L, Hannon GJ (2004) Argonaute2 is the catalytic engine of mammalian RNAi. Science 305: 1437–1441

Liu Y, Teng C, Xia R, Meyers BC (2020) PhasiRNAs in Plants: Their Biogenesis, Genic Sources, and Roles in Stress Responses, Development, and Reproduction. The Plant cell 32: 3059–3080

Martin M (2011) Cutadapt removes adapter sequences from high-throughput sequencing reads. EMBnetjournal; Vol 17, No 1: Next Generation Sequencing Data Analysis

Martinez de Alba AE, Moreno AB, Gabriel M, Mallory AC, Christ A, Bounon R, Balzergue S, Aubourg S, Gautheret D, Crespi MD et al (2015) In plants, decapping prevents RDR6-dependent production of small interfering RNAs from endogenous mRNAs. Nucleic acids research 43: 2902–2913

Meister G (2013) Argonaute proteins: functional insights and emerging roles. Nature reviews Genetics 14: 447–459

Meister G, Landthaler M, Patkaniowska A, Dorsett Y, Teng G, Tuschl T (2004) Human Argonaute2 mediates RNA cleavage targeted by miRNAs and siRNAs. Molecular cell 15: 185–197

Motamedi MR, Verdel A, Colmenares SU, Gerber SA, Gygi SP, Moazed D (2004) Two RNAi complexes, RITS and RDRC, physically interact and localize to noncoding centromeric RNAs. Cell 119: 789–802

Mourrain P, Beclin C, Elmayan T, Feuerbach F, Godon C, Morel J-B, Jouette D, Lacombe A-M, Nikic S, Picault N et al (2000) *Arabidopsis SGS2* and *SGS3* genes are required for posttranscriptional gene silencing and natural virus resistance. Cell 101: 533–542

Nagano H, Fukudome A, Hiraguri A, Moriyama H, Fukuhara T (2014) Distinct substrate specificities of Arabidopsis DCL3 and DCL4. Nucleic acids research 42: 1845–1856

Nielsen CPS, Arribas-Hernández L, Han L, Reichel M, Woessmann J, Daucke R, Bressendorff S, López-Márquez D, Andersen SU, Pumplin N et al (2024) Evidence for an RNAi-independent role of Arabidopsis DICER-LIKE2 in growth inhibition and basal antiviral resistance. The Plant cell

Nour-Eldin HH, Hansen BG, Norholm MH, Jensen JK, Halkier BA (2006) Advancing uracil-excision based cloning towards an ideal technique for cloning PCR fragments. Nucleic acids research 34: e122

Orban TI, Izaurralde E (2005) Decay of mRNAs targeted by RISC requires XRN1, the Ski complex, and the exosome. Rna 11: 459–469

Pastore B, Hertz HL, Price IF, Tang W (2021) pre-piRNA trimming and 2’-O-methylation protect piRNAs from 3’-tailing and degradation in *C. elegans*. Cell reports 36

Peragine A, Yoshikawa M, Wu G, Albrecht HL, Poethig RS (2004) SGS3 and SGS2/SDE1/RDR6 are required for juvenile development and the production of trans-acting siRNAs in Arabidopsis. Genes & development 18: 2368–2379

Poulsen C, Vaucheret H, Brodersen P (2013) Lessons on RNA silencing mechanisms in plants from eukaryotic argonaute structures. The Plant cell 25: 22–37

Ren G, Xie M, Zhang S, Vinovskis C, Chen X, Yu B (2014) Methylation protects microRNAs from an AGO1-associated activity that uridylates 5’ RNA fragments generated by AGO1 cleavage. Proceedings of the National Academy of Sciences of the United States of America 111: 6365–6370

Ren Y, Ma X, Song B, Yang X, Chen Y, Yu Y, Chen X, Mo B, Wang X (2023) HEN1 SUPPRESSOR1 stabilizes polymerase IV RNAs via uridylation in Arabidopsis. Plant physiology 193: 186–189

Rosso MG, Li Y, Strizhov N, Reiss B, Dekker K, Weisshaar B (2003) An Arabidopsis thaliana T-DNA mutagenized population (GABI-Kat) for flanking sequence tag-based reverse genetics. Plant molecular biology 53: 247–259

Saito K, Sakaguchi Y, Suzuki T, Suzuki T, Siomi H, Siomi MC (2007) Pimet, the Drosophila homolog of HEN1, mediates 2′-O-methylation of Piwi-interacting RNAs at their 3′ ends. Genes & development 21: 1603–1608

Sakurai Y, Baeg K, Lam AYW, Shoji K, Tomari Y, Iwakawa H-o (2021) Cell-free reconstitution reveals the molecular mechanisms for the initiation of secondary siRNA biogenesis in plants. Proceedings of the National Academy of Sciences 118: e2102889118

Scheer H, de Almeida C, Ferrier E, Simonnot Q, Poirier L, Pflieger D, Sement FM, Koechler S, Piermaria C, Krawczyk P et al (2021) The TUTase URT1 connects decapping activators and prevents the accumulation of excessively deadenylated mRNAs to avoid siRNA biogenesis. Nature communications 12: 1298

Schiebel W, Haas B, Marinković S, Klanner A, Sänger HL (1993a) RNA-directed RNA polymerase from tomato leaves. I. Purification and physical properties. Journal of Biological Chemistry 268: 11851–11857

Schiebel W, Haas B, Marinković S, Klanner A, Sänger HL (1993b) RNA-directed RNA polymerase from tomato leaves. II. Catalytic in vitro properties. Journal of Biological Chemistry 268: 11858–11867

Schiebel W, Pelissier T, Reidel L, Thalmeir S, Schiebel R, Kempe D, Lottspeich F, Sanger HL, Wassenegger M (1998) Isolation of an RNA-directed RNA polymerase-specific cDNA clone from tomato. The Plant cell 10: 2087–2102

Sement FM, Ferrier E, Zuber H, Merret R, Alioua M, Deragon JM, Bousquet-Antonelli C, Lange H, Gagliardi D (2013) Uridylation prevents 3’ trimming of oligoadenylated mRNAs. Nucleic acids research 41: 7115–7127

Shen B, Goodman HM (2004) Uridine addition after microRNA-directed cleavage. Science 306: 997

Shukla A, Yan J, Pagano DJ, Dodson AE, Fei Y, Gorham J, Seidman JG, Wickens M, Kennedy S (2020) poly(UG)-tailed RNAs in genome protection and epigenetic inheritance. Nature 582: 283–288

Si-Ammour A, Windels D, Arn-Bouldoires E, Kutter C, Ailhas J, Meins F, Jr., Vazquez F (2011) miR393 and Secondary siRNAs Regulate Expression of the TIR1/AFB2 Auxin Receptor Clade and Auxin-Related Development of Arabidopsis Leaves Plant physiology 157: 683–691

Song J, Wang X, Song B, Gao L, Mo X, Yue L, Yang H, Lu J, Ren G, Mo B et al (2019) Prevalent cytidylation and uridylation of precursor miRNAs in Arabidopsis. Nature Plants 5: 1260–1272

Song JJ, Smith SK, Hannon GJ, Joshua-Tor L (2004) Crystal structure of Argonaute and its implications for RISC slicer activity. Science 305: 1434–1437

Song M-S, Rossi JJ (2017) Molecular mechanisms of Dicer: endonuclease and enzymatic activity. Biochemical Journal 474: 1603–1618

Tsai H-Y, Chen C-Chieh G, Conte D, Jr., Moresco James J, Chaves Daniel A, Mitani S, Yates John R, III, Tsai M-D, Mello Craig C (2015) A Ribonuclease Coordinates siRNA Amplification and mRNA Cleavage during RNAi. Cell 160: 407–419

Tsuboi T, Kuroha K, Kudo K, Makino S, Inoue E, Kashima I, Inada T (2012) Dom34:Hbs1 Plays a General Role in Quality-Control Systems by Dissociation of a Stalled Ribosome at the 3&#x2032; End of Aberrant mRNA. Molecular cell 46: 518–529

Tsutsui H, Higashiyama T (2017) pKAMA-ITACHI Vectors for Highly Efficient CRISPR/Cas9-Mediated Gene Knockout in Arabidopsis thaliana. Plant and Cell Physiology 58: 46–56

Tu B, Liu L, Xu C, Zhai J, Li S, Lopez MA, Zhao Y, Yu Y, Ramachandran V, Ren G et al (2015a) Distinct and Cooperative Activities of HESO1 and URT1 Nucleotidyl Transferases in MicroRNA Turnover in Arabidopsis. PLoS genetics 11: e1005119

Tu B, Liu L, Xu C, Zhai J, Li S, Lopez MA, Zhao Y, Yu Y, Ramachandran V, Ren G et al (2015b) Distinct and Cooperative Activities of HESO1 and URT1 Nucleotidyl Transferases in MicroRNA Turnover in Arabidopsis. PLoS genetics 11: e1005119

Vermeulen A, Behlen L, Reynolds A, Wolfson A, Marshall WS, Karpilow J, Khvorova A (2005) The contributions of dsRNA structure to Dicer specificity and efficiency. Rna 11: 674–682

Vigh ML, Bressendorff S, Thieffry A, Arribas-Hernández L, Brodersen P (2022) Nuclear and cytoplasmic RNA exosomes and PELOTA1 prevent miRNA-induced secondary siRNA production in Arabidopsis. Nucleic acids research 50: 1396–1415

Wang X, Kong W, Wang Y, Wang J, Zhong L, Lao K, Dong X, Zhang D, Huang H, Mo B et al (2022) Uridylation and the SKI complex orchestrate the Calvin cycle of photosynthesis through RNA surveillance of TKL1 in Arabidopsis. Proceedings of the National Academy of Sciences 119: e2205842119

Wang X, Wang Y, Dou Y, Chen L, Wang J, Jiang N, Guo C, Yao Q, Wang C, Liu L et al (2018) Degradation of unmethylated miRNA/miRNA*s by a DEDDy-type 3’ to 5’ exoribonuclease Atrimmer 2 in *Arabidopsis*. Proceedings of the National Academy of Sciences 115: E6659–E6667

Wang X, Zhang S, Dou Y, Zhang C, Chen X, Yu B, Ren G (2015) Synergistic and Independent Actions of Multiple Terminal Nucleotidyl Transferases in the 3’ Tailing of Small RNAs in Arabidopsis. PLoS genetics 11: e1005091

Xie Z, Allen E, Wilken A, Carrington JC (2005) DICER-LIKE 4 functions in trans-acting small interfering RNA biogenesis and vegetative phase change in Arabidopsis thaliana. Proceedings of the National Academy of Sciences of the United States of America 102: 12984–12989

Yoshikawa M, Han Y-W, Fujii H, Aizawa S, Nishino T, Ishikawa M (2021) Cooperative recruitment of RDR6 by SGS3 and SDE5 during small interfering RNA amplification in Arabidopsis. Proceedings of the National Academy of Sciences 118: e2102885118

Yoshikawa M, Iki T, Numa H, Miyashita K, Meshi T, Ishikawa M (2016) A Short Open Reading Frame Encompassing the MicroRNA173 Target Site Plays a Role in trans-Acting Small Interfering RNA Biogenesis. Plant physiology 171: 359–368

Yoshikawa M, Iki T, Tsutsui Y, Miyashita K, Poethig RS, Habu Y, Ishikawa M (2013) 3’ fragment of miR173-programmed RISC-cleaved RNA is protected from degradation in a complex with RISC and SGS3. Proceedings of the National Academy of Sciences of the United States of America 110: 4117–4122

Yoshikawa M, Peragine A, Park MY, Poethig RS (2005) A pathway for the biogenesis of trans-acting siRNAs in Arabidopsis. Genes & development 19: 2164–2175

Yu B, Yang Z, Li J, Minakhina S, Yang M, Padgett RW, Steward R, Chen X (2005) Methylation as a crucial step in plant microRNA biogenesis. Science 307: 932–935

Zhang C, Ng DW-K, Lu J, Chen ZJ (2012) Roles of target site location and sequence complementarity in trans-acting siRNA formation in Arabidopsis. The Plant Journal 69: 217–226

Zhang X, Zhu Y, Liu X, Hong X, Xu Y, Zhu P, Shen Y, Wu H, Ji Y, Wen X et al (2015) Suppression of endogenous gene silencing by bidirectional cytoplasmic RNA decay in Arabidopsis. Science 348: 120–123

Zhang Z, Hu F, Sung MW, Shu C, Castillo-González C, Koiwa H, Tang G, Dickman M, Li P, Zhang X (2017) RISC-interacting clearing 3’-5’ exoribonucleases (RICEs) degrade uridylated cleavage fragments to maintain functional RISC in Arabidopsis thaliana. eLife 6: e24466

Zuber H, Scheer H, Ferrier E, Sement FM, Mercier P, Stupfler B, Gagliardi D (2016) Uridylation and PABP Cooperate to Repair mRNA Deadenylated Ends in Arabidopsis. Cell reports 14: 2707–2717

Zuber H, Scheer H, Joly AC, Gagliardi D (2018) Respective Contributions of URT1 and HESO1 to the Uridylation of 5’ Fragments Produced From RISC-Cleaved mRNAs. Front Plant Sci 9: 1438

